# IDH-Tau-EGFR triad defines the neovascular landscape of diffuse gliomas by controlling mesenchymal differentiation

**DOI:** 10.1101/541326

**Authors:** Ricardo Gargini, Berta Segura-Collar, Beatriz Herránz, Vega Garcia-Escudero, Andrés Romero-Bravo, Felipe J Núñez, Daniel García-Pérez, Jacqueline Gutiérrez-Guamán, Angel Ayuso-Sacido, Joan Seoane, Angel Pérez-Núñez, Juan M. Sepúlveda-Sánchez, Aurelio Hernández-Laín, María G. Castro, Ramón García-Escudero, Jesús Ávila, Pilar Sánchez-Gómez

**Author notes:** Equal contribution (Ricardo Gargini, Berta Segura-Collar). Correspondence to, Dr. Pilar Sanchez-Gomez, Neurooncology Unit, Instituto de Salud Carlos III-UFIEC, Madrid, Spain, Crtra/ Majadahonda-Pozuelo, Km 2, Majadahonda, 28220, Spain. Correspondence to, Prof. Jesús Ávila, Department of Neurobiology, Centro de Biología Molecular Severo Ochoa, Cantoblanco 28049, C/Nicolás Cabrera 1, 28049 Madrid, Spain.

## Abstract

Mutant IDH1/2 gliomas represent a more indolent form of cancer. However, how this group of tumors evolve, in a microenvironment-dependent manner, is still a pending question. Here we describe that the expression of *Tau*, a gene classically associated with neurodegenerative diseases, is epigenetically controlled by the balance between wild-type and mutant IDH1/2 in gliomas. Moreover, the level of *Tau* decreases when the tumor progresses. Besides, Tau is almost absent from tumors with EGFR mutations, whereas its expression is inversely correlated with overall survival in gliomas carrying wild-type or amplified EGFR. Here, we demonstrate that the overexpression of Tau, through the stabilization of microtubules, impairs the mesenchymal/pericyte-like transformation of glioma cells by blocking the EGFR-NFκB-TAZ axis. However, mutant EGFR induces a constitutive activation of this pathway, which is no longer sensitive to Tau. By inhibiting the phenotypic plasticity of EGFRamp/wt glioma cells, Tau protein inhibits angiogenesis and favors vascular normalization, decreasing tumor aggressiveness and rendering the tumors more sensitive to chemotherapy.

**One Sentence Summary:** Tau, which is induced by IDH mutations, inhibits the EGFR/NF-kB/TAZ axis and impairs the mesenchymal/pericyte-like transformation of glioma cells, normalizing the vasculature and impairing tumor aggressiveness.

## INTRODUCTION

Diffuse gliomas are classified and graded according to histological criteria. They include low and intermediate-grade gliomas (herein called Lower-Grade Gliomas, LGG), which encompass World Health Organization (WHO) grades 2 and 3, and the highly aggressive WHO grade 4 glioblastomas (GBM), with 5-year survival rates of 5%. They are categorized based on the increments in cellular atypia and mitotic activity. On top of that, GBM are characterized by the specific presence of areas of necrosis and robust neoangiogenesis, being considered one of the most vascularized cancers. LGG are more indolent tumors although many of them progress into secondary GBM, albeit at highly variable intervals, with survival rates that goes from 1 to 15 years (1). Unfortunately, little is known about the factors that drive this transition from LGG to GBM.

A big effort has been made in the last two decades in order to characterize the genetic modifications associated with gliomas. Some of them have been incorporated into the novel WHO classification. Particularly, the identification of mutations in the *IDH1/2* (*Isocitrate dehydrogenase 1/2*) genes, which are associated with a more favorable prognosis in gliomas (2), is common now in the routine clinical practice. IDH1/2 mutated proteins induce the accumulation of the oncometabolite, 2-D-hydroxyglutarate (2-HG), which competes with the α-ketoglutarate (α-KG) produced by the wild-type IDH enzymes and blocks TET (Ten Eleven Translocation)-mediated DNA demethylation. This process generates a CpG island methylator phenotype (G-CIMP), which is associated with a general suppression of gene expression (3). Moreover, 2-HG inhibits histone demethylases, which further contribute to this phenotype (4). By contrast, wild-type IDH1 promotes the metabolic adaptation of GBM cells to support aggressive growth (5). Therefore, it has been proposed that the balance between wild-type and mutant IDH1/2 function determines the clinical outcome of gliomas, including their sensitivity to radiation and chemotherapy (6).

Within the new subclasses of high grade gliomas (proneural (PN), classic (CL) or mesenchymal (MES)), *IDH1/2* mutations are accumulated in the first group, which is enriched in secondary GBM and includes tumors with a better clinical prognosis (7). By contrast, mutations in *EGFR* (Epidermal growth factor receptor) accumulate in the CL and MES subtypes. This gene is mutated and/or amplified in a large percentage of diffuse gliomas and it has been associated with proliferation and survival, as well as with the invasive properties of glioma cells (7;8).

Several cytoskeletal proteins have been involved in tumor progression. Tau, encoded by the gene *MAPT* (Microtubule-associated protein tau), is well-known for its relevance in Alzheimer’s disease (AD) although it is also expressed in healthy brains, where it controls neural development and synaptic transmission (9). In addition, Tau and other microtubule-stabilizing agents like taxanes modulate protein and organelle trafficking (10,11), which could be relevant for cancer cells. Interestingly, a possible co-morbidity of dementias and GBM had been suggested (12), which led us to perform a bioinformatic analysis. We found that *Tau (MAPT)*, among other genes related to neurodegeneration, is expressed in gliomas, where it seems to correlate negatively with tumor progression (Gargini et al., Front. Aging Neurosci., in press). Based on these data, we decided to conduct a more comprehensive characterization of Tau in this deadly pathology. Here we show that the expression of this protein depends on the genetic status of *IDH1/2*, being enriched in LGG and PN gliomas, where it hinders tumor progression. Mechanistically, we have found that Tau inhibits the EGFR-NF-kB-TAZ signaling pathway, provided that no EGFR mutations are present. By blocking this cascade, it impedes the plasticity of the tumor cells and their capacity to generate mesenchymal/pericyte-like cells, which participate in the processes of angiogenesis and neo-vascularization. As a consequence, Tau favors the normalization of the glioma’s vasculature and hampers tumor progression. Therefore, we propose a novel role for Tau in gliomas, acting downstream of IDH mutations to orchestrate the vascular phenotype and the aggressiviness of these tumors.

## RESULTS

### High levels of Tau (MAPT) correlate inversely with glioma aggressiveness

The in silico analysis of the glioma TCGA data set showed that the level of *Tau (MAPT)* decreases as the tumor grade increases, at least in astrocytomas (Fig. 1A, and fig. S1 A and B). In agreement with this observation, a higher expression of the *Tau (MAPT)* gene was associated with an increased overall survival of glioma patients (Fig. 1B and C, fig S1C to F). These results confirmed our previous data (Gargini et al., Front. Aging Neurosci., in press) and prompted us to perform an immunohistochemical (IHC) staining on glioma samples, which showed that Tau protein is clearly expressed in the cytoplasm of tumor cells, with a very different pattern to the one observed in normal tissue (NT) (Fig. 1D, and fig. S1G). Moreover, we found high levels of Tau in a subset of the gliomas analyzed by Western Blot (WB) (Fig. 1E). The quantification of the IHC staining (Fig. 1F) and the WB (Fig. 1G) confirmed that Tau protein is clearly enriched in LGG compared to GBM. As this accumulation could explain by itself the survival data (Fig. 1B and C), we decided to dissect out the effect of *Tau* expression in LGG and GBM by using separated TCGA cohorts. Fig. 2A and B show that the transcript level of *Tau* correlates with an increased overall suvival in both sets of patients. Similar results were obtained when we meassured *Tau* by RT-PCR analysis in our own GBM cohort (Fig. 2C). Collectively, these results support the idea that *Tau* expression is associated with a less aggressive behavior of gliomas, independently of the tumor grade. Furthermore, we found a marked decrease in the expression of Tau in disease free vs progressed tumors (Fig. 2D). To confirm these observations we performed a longitudinal IHC quantification of Tau in primary LGG and their recurrent paired samples. Remarkably, we found a consistent decrease in Tau expression in those tumors that had progress into a more aggressive phenotype (Fig. 2E), which was not observed when there was no change in the histological classification of the recidives (Fig. 2F). These results suggest that Tau downregulation must be important for the tumors to relapse after surgical resection, which is a critical step in the mortality related to this pathology.

**Fig. 1.**
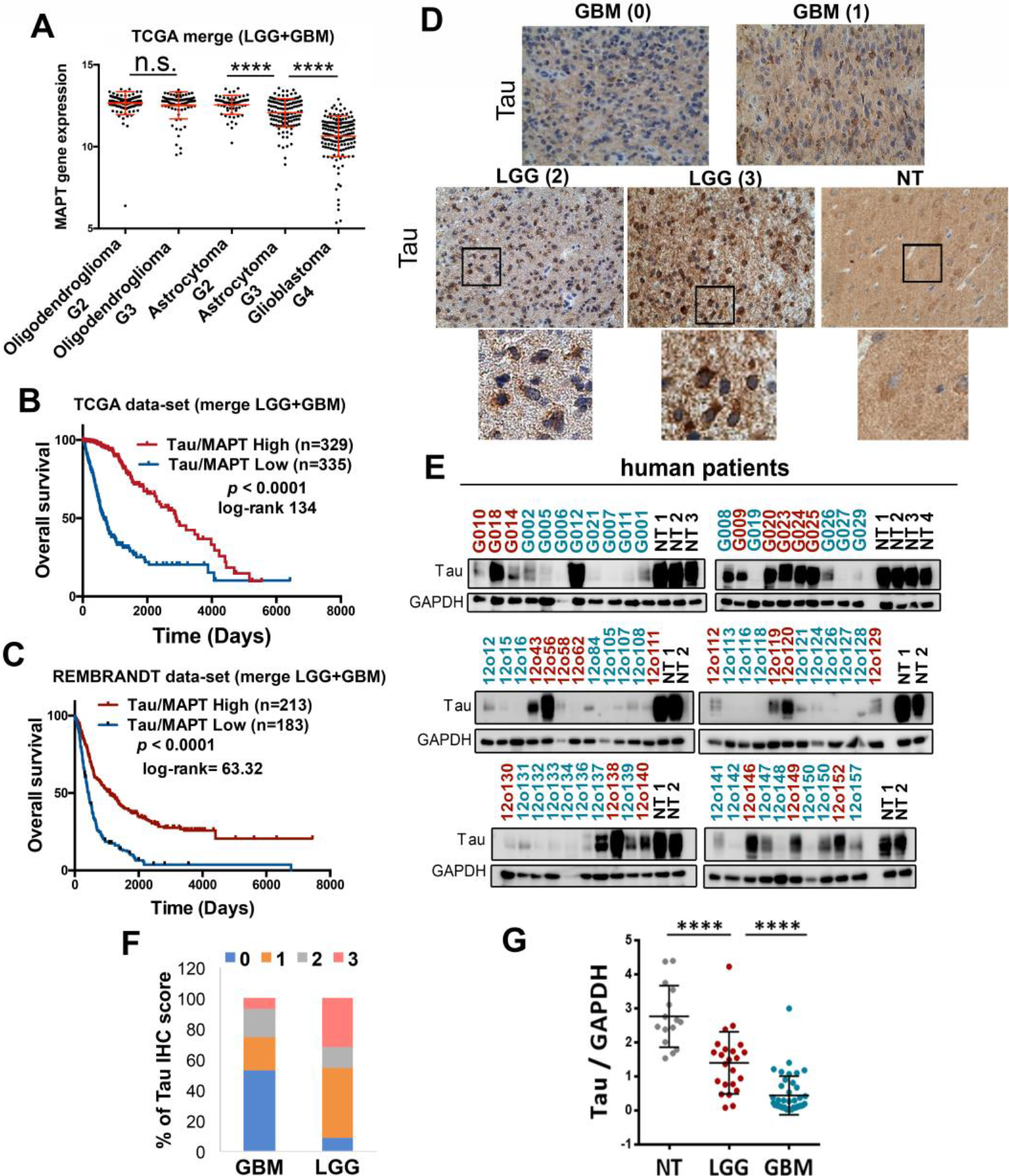
Tau is expressed in gliomas and it is enriched in lower-grade tumors. (**A**) Analysis of *Tau (MAPT)* mRNA expression by RNAseq in gliomas (TCGA cohort) grouped according to the WHO classification (histological type and grade) (n=692). (**B** and **C**) Kaplan-Meier overall survival curves of patients from the TCGA (LGG+GBM) (n=664) and the Rembrandt (LGG+GBM) (n=396) cohorts. Patients in each cohort were stratified into 2 groups based on high and low *Tau (MAPT)* expression values. (**D**) Representative pictures of the IHC Tau staining of several gliomas and normal tissue (NT). The Tau IHC score is represented between brackets and an amplified section of the last three images is shown on the bottom. (**E**) WB analysis of Tau expression in tumor tissue extracts from patients diagnosed with LGG (red) and GBM (blue). NT was used as a control of Tau expression and GAPDH level as a loading control. (**F**) Percentage of tumors (GBM (n=55) and LGG (n=22)) with different Tau IHC score. (**G**) Quantification of the relative amount of Tau in the WB in E. ****, *p* ≤0.0001. n.s. non significant.

**Fig. 2.**
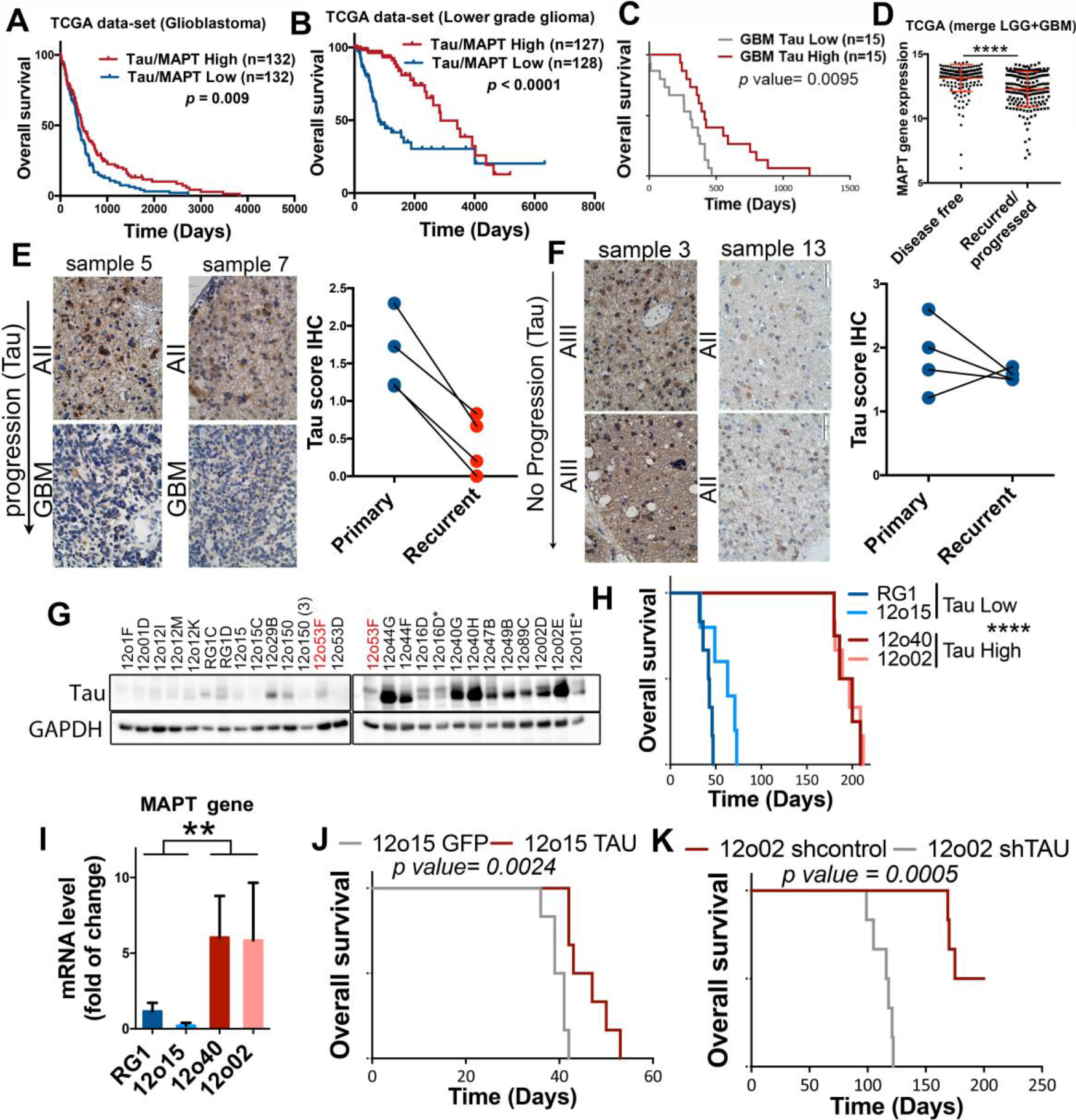
The expression of Tau in gliomas correlates inversely with tumor aggressiveness. (**A** and **B**) Kaplan-Meier overall survival curves of patients from the TCGA, the GBM cohort (n=264) (A) or the LGG cohort (n= 155). Patients in each cohort were stratified into 2 groups based on high and low *Tau (MAPT)* expression values. (**C**) Kaplan-Meier overall survival curves of patients from our own GBM cohort (n=30). Patients in each cohort were stratified into 2 groups based on high and low *Tau (MAPT)* expression values (measured by RT-PCR using *HPRT* levels for normalization). (**D**) Analysis of *Tau (MAPT)* mRNA expression by RNAseq in gliomas (TCGA cohort) grouped according to the clinical evolution of the tumors (n=386). (**E** and **F**) Representative pictures of the IHC Tau staining of paired glioma samples (primary and recurrent tumor). Average Tau IHC score of the paired samples in shown on the right. (**G**) WB analysis of Tau expression in tumor tissue extracts from subcutaneous patient-derived xenografts (PDXs). The same extract from 12o53F cells was loaded in both gels for comparison.GAPDH was used as a loading control. (**H**) Kaplan-Meier overall survival curves of mice that were orthotopically injected with different primary GBM cell lines. (**I**) qRT-PCR analysis of *Tau (MAPT)* expression in tumor tissue extracts from (H). HPRT expression was used for normalization. (**J**) Kaplan-Meier overall survival curves of mice that were orthotopically injected with 12o15 cells overexpressing GFP or Tau (n=6). (**K**) Kaplan-Meier overall survival curves of mice that were orthotopically injected with 12o02 control or 12o02-shTau cells (n=6). The statistical analysis is shown on the bottom. ****, *p* ≤0.0001.

To further asses the presence of Tau in glioma cells and its participation in tumor aggressiveness we analyzed its expression in a panel of patient-derived-xenografts (PDX), obtained from GBM surgeries and grown subcutaneously, devoided of any trapped neurons. As we had previously observed in tumor samples (Fig. 1E), the amount of Tau protein in the PDXs was highly variable (Fig. 2G, and fig. S2A). It is important to remark that in those tumors with high levels of the protein, a big percentage of GFAP cells were positive for Tau staining (fig S2A and fig. S2B), further supporting the specific expression of Tau in glioma cells.

In order to know if Tau levels affect tumor behavior, we injected some of the PDX cells into the brains of immunodeficient mice. We observed that those primary cell lines with higher levels of Tau (measured by qRT-PCR with human specific primers (Fig. 2I) grew significantly slowlier (Fig. 2H). Moreover, the overexpression of this gene in a Tau-deficient glioma cell line (12o15) delayed tumor formation (Fig. 2J and fig. S2C), whereas its downregulation in Tau-enriched cells (12o02) clearly increased their aggressive behavior (Fig. 2K and fig. S2D). Altogether, our results confirm that Tau is expressed in the tumor cells of several gliomas, especially in the less aggressive ones. Additionally, they suggest that Tau could be playing an active role as an inhibitor of tumor progression.

### The expression of Tau (MAPT) in gliomas is regulated by IDH function

We analyzed the genetic background of gliomas in relation to the levels of *Tau (MAPT)* and we found that *IDH1* mutations accumulate in high-*Tau* gliomas (Fig. 3A). This correlation was validated using a Volcano Plot analysis (Fig. 3B). Furthermore, Tau protein was detected in the majority of tumor cells that express the most common IDH1 mutation (R132H) (fig. S3A and B). This observation confirms that the protein is truly expressed in the tumor cells and not only in the trapped neurons. We then quantified the amount of Tau IHC staining in wild-type and mutant *IDH* gliomas and we found a clear enrichment of high and medium stained sections in the second group (Fig. 3C). These results suggest that there is a strong correlation between the presence of IDH mutations and the expression of Tau. This seems to be independent of the tumor grade as we found higher levels of Tau in *IDH* mutant compared to *IDH* wild-type tumors in the GBM (fig. S3C), as well as in the LGG (fig. S3D) TCGA cohorts. In order to confirm the relationship between Tau and IDH mut, we analyzed a recently published glioma model, in which IDH1 wt or IDH R132H had been expressed in an *ATRX* mutant background (13). As expected, there was a clear delay in the tumor growth after the expression of mutant IDH1 (Fig. 3D). Moreover, when we dissected out the intracranial tumors we measured a clear increase in Tau protein levels in the mutant compared to the wild-type allografts (Fig. 3E). These results suggest that the expression of Tau in gliomas is induced by mutant IDH proteins.

**Fig. 3.**
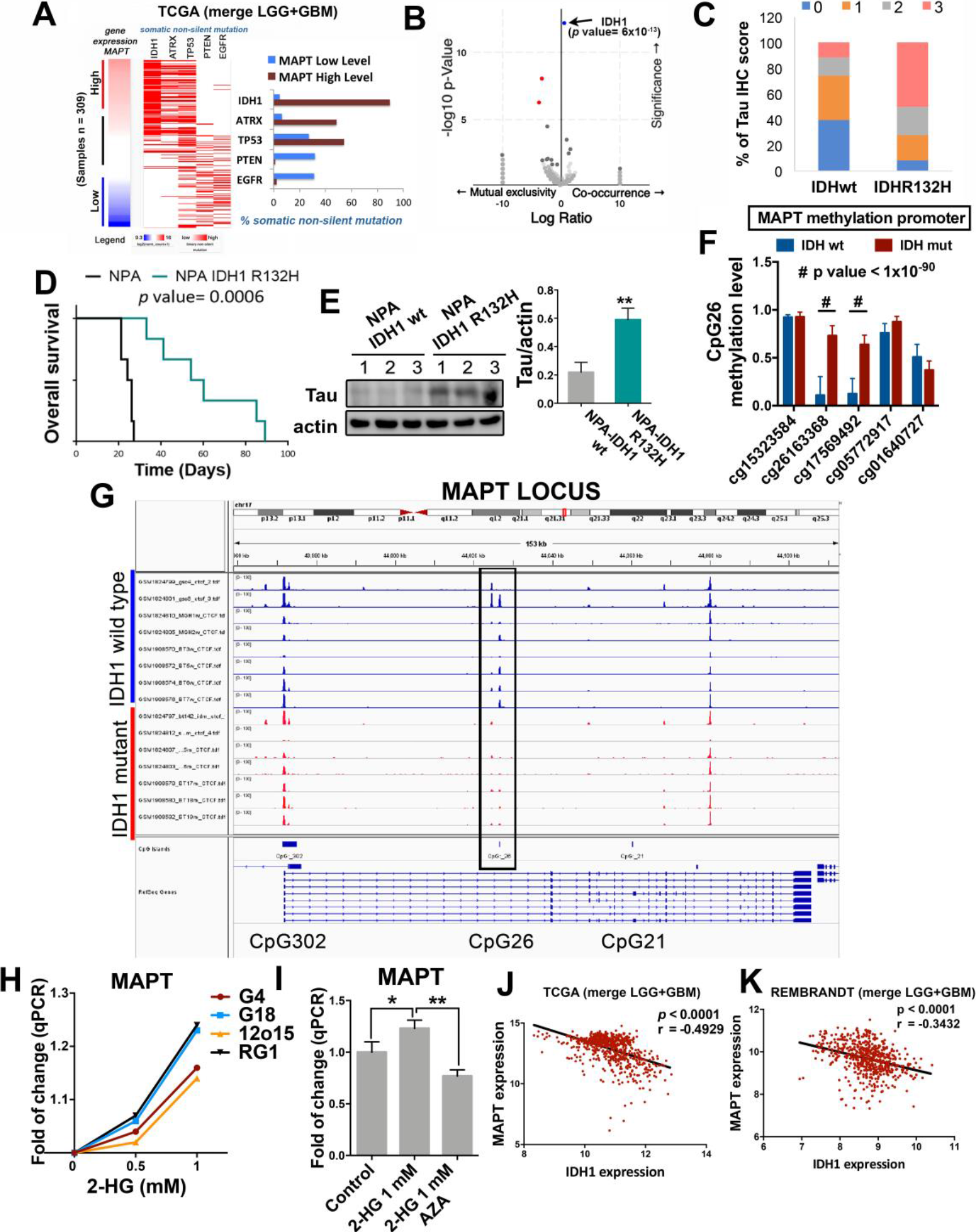
*Tau* expression is regulated by IDH1/2 function. (**A**) Analysis of non-silent somatic mutations in genes commonly modified in diffuse glioma grouped based on high or low expression of *Tau (MAPT)*. (**B**) Volcano plots showing mutated genes with differential distribution in glioma comparing tumors with high and low level of *Tau*. The arrow points to *IDH1* mutations. (**C**) Percentage of tumors with different Tau IHC score in wt (n= 35) and mutant (n=36) *IDH1* gliomas. (**D**) Kaplan-Meier overall survival curves of mice that were orthotopically injected with NPA IDH1 wt or NPA-IDH1 R132H cells (n=6). (**E**) WB analysis and quantification of Tau expression in intracranial tumors from (D). Actin was used as loading control. (**F**) Quantification of the methylation of CpG26 using 5 different probes and comparing IDHwt vs IDHmut gliomas. (**G**) CTCF-binding profiles for the *Tau (MAPT*) CpG26 locus in IDH-mutant and IDH wild-type tumors, normalized by average signal. (**H**) Analysis of the expression of *MAPT* gene expression by qPCR in the presence of increasing amounts of 2-hydroxy-glutarate (2-HG) in RG1, 12o15, GB4 and GB18 cells. (**I**) Analysis of the expression of *MAPT* gene expression by qRT-PCR in RG1 cells cultured in the presence of 1 mM of the 2-HG, with or without azacytidine (AZA) (1 μM). (**J** and **K**) Correlation of the expression of *Tau (MAPT)* with that of wild-type *IDH1* using the TCGA-merge (LGG+GBM) (n=) (J) and the Rembrandt (LGG+GBM) (n=580) (K) cohorts. **, *p* ≤0.01; ****, *p* ≤0.0001; #, *p* ≤1×10^−90^

The G-CIMP phenotype, which is associated with the presence of *IDH* mutations, is supposed to block the transcription of many genes. However, it also activates certain others, especially those involved in the tumorigenesis of LGG (like the *PDGFRA (Platelet Derived Growth Factor Receptor Alpha)* oncogene) through the disruption of the repressive structure of the CTCF insulator protein (14). Interestingly, we observed an increased expression of *Tau* in the G-CIMP GBM subtype (fig. S4A), as well as a significant correlation between the amount of *Tau (MAPT)* and *PDGFRA* mRNAs in the TCGA samples (fig. S4B). These results suggest that Tau promoter might be controlled by the epigenetic changes induced by IDH mutations. To test this theory, we first analyzed the presence of CpG islands in the *Tau (MAPT*) promoter region by using the xena genome browser. We identified three of them in the 5′region of transcription initiation site (fig. S4C), which correlate with the sites previously described (15). Using epigenetic data from the TCGA (LGG cohort) we compared the methylation of the whole promoter region in mutant vs wild-type IDH tumors. We observed an increased methylation in CpG:26 (Fig. 3F, fig. S4D and E) and in some parts of CpG:302 (fig. S4D and E) in the mutant tumors. However, when we analyzed the ChIP-seq data of the CTCF binding to the different clusters, we observed that only the binding to the CpG:26 site was lost in the *IDHmut* tumors (Fig. 3G). In order to obtain an independent confirmation of the epigenetic upregulation of *Tau (MAPT)* in response to mutant IDH, we treated primary GB cells with 2-HG and we observed a dose-dependent accumulation of its mRNA (Fig. 3H), an effect that was reversed in the presence of azacytidine (AZA) (Fig. 3I). In summary, these results strongly suggest that the increase in the methylation of the CpG:26 region, induced by IDH mutant proteins, changes the chromosomal insulator topology and the binding of CTCF to the *Tau* promoter, activating its transcription.

It is well known that *IDH1/2* mutations define a distinct subset of gliomas with a better outcome. By contrast the expression of *IDH* wild-type in LGG and GBM defines a subgroup with poorer prognosis (fig. S4F). Interestingly, we found an inverse correlation between the expression of wild-type *IDH1* and *Tau* on the TCGA (Fig. 3J) and the Rembrandt (Fig. 3K) cohorts, suggesting that the balance between wild-type and mutant IDH expression determines the epigenetic regulation of the *Tau* promoter.

### Tau opposes EGFR signaling in gliomas

The results presented here indicate that Tau is highly expressed in a subset of gliomas, especially in *IDH* mutant tumors, where it seems to impair tumor aggressiveness. To gain insight into the mechanism behind this behavior, we performed a DAVID gene analysis of the pathways co-upregulated with *Tau* in gliomas. As expected, we found a positive association with microtubule and neurogenic-related processes. However, the other Tau-linked pathways (vesicle-transport, MAPK/GTPase regulation and Pleckstrin-related processes) were somehow related to receptor tyrosine kinases signaling (fig. S5A). In addition, our in silico analysis indicated that *IDH* mutations, as well as higher levels of *Tau* are mutually exclusive with *EGFR* and *PTEN* mutations (Fig. 3A, and Fig. 4A, and fig. S5B), which was confirmed in a Volcano Plot analysis (Fig. 4B). To interrogate if Tau could be modulating this signaling pathway, we expressed IDH wt or IDH R132H in a EGFR amplified cell line (RG1). As we have previously observed in the mouse glioma model (Fig. 3D), IDH mut impaired tumor burden (Fig. 4C). However, we also confirmed that the overexpression of IDHwt increases the aggressiveness of the glioma cells (Fig. 4C). These effects were paralleled by changes in Tau expression, which augmented in IDH mut and decreased in IDH wt tumors, compared to the control ones (Fig. 4D and E). This result is in agreement with the in silico data (Fig. 3J an K), and allows us to propose that the induction of Tau transcription might depend on opposite functions between wild-type and mutant IDH expression. Interestingly, the phosphorylation of EGFR (phosphor-EGFR) showed a striking inverse correlation with Tau levels in the mouse tumors (Fig. 4D and F), as well as in human samples (fig. S5C and D). More importantly, the expression of *Tau* correlated with overall survival in the group of *EGFR*amp/wt gliomas (Fig. 4G) but it had no clinical relevance in the presence of additional mutations in *EGFR* (Fig. 4H). Taken together, these results suggest that there could be a negative effect of IDH/Tau on the pathway activated by this receptor, but only in the group of *EGFR*amp/wt gliomas.

**Fig. 4.**
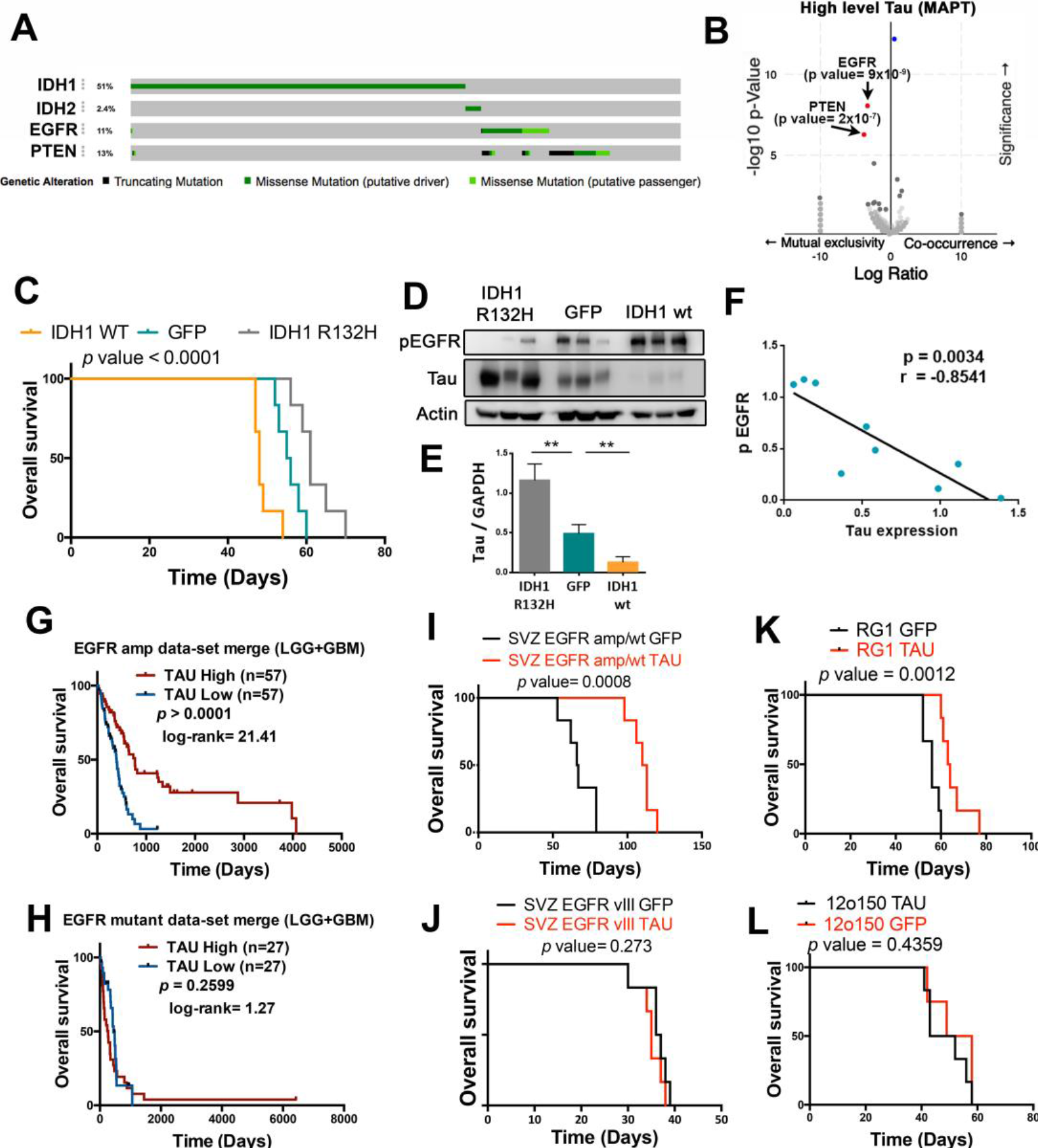
Tau opposes EGFR in gliomas. (**A**) Distribution of somatic non-silent mutations in *IDH1/2*, *EGFR* and *PTEN* in a glioma cohort (TCGA, n=812). (**B**) Volcano plots showing mutated genes with differential distribution in gliomas, comparing tumors with high and low level of *Tau*. The arrows point to *PTEN* and *EGFR* mutations. (**C**) Kaplan-Meier overall survival curves of mice that were orthotopically injected with RG1, RG1 IDH1 wt or RG1 IDH1 R32H cells (n=6). (**D**) WB analysis of phosphorylated EGFR (pEGFR) and Tau in intracranial tumors from (C). Actin was used as a loading control. (**E**) Quantification of the amount of Tau in (D). (**F**) Correlation between the levels of Tau and phospho-EGFR in RG1 tumors. (**G** and **H**) Kaplan-Meier overall survival curves of patients from the LGG+GBM TCGA cohort. Patients were separated based on the *EGFR* status: tumors without mutations (amplified or wild type) (n=114) (G) and amplified tumors with mutations (n=54) (H). Then, they were stratified into 2 groups based on *Tau (MAPT)* expression value. (**I** to **L)** Kaplan-Meier overall survival curves of mice that were orthotopically injected with SVZ-EGFRamp/wt (I), SVZ-EGFRvIII (J), RG1 (*EGFR*amp) (K) or 12o150 (*EGFR*mut) (L) cells, overexpressing either GFP or Tau (n=6). **, *p* ≤ 0.01.

To evaluate the effect of Tau on EGFR signaling, we overexpressed this protein in two mouse glioma models: SVZ-EGFRamp/wt and SVZ-EGFRvIII, and we analized their capacity to grow as intracranial allografts. These models were generated by transforming subventricular zone (SVZ) progenitors from p16/p19 ko mice with retrovirus expressing either the wild-type or the variant III (vIII) isoform of the receptor. Both type of cells depend on EGFR signaling *in vitro* and *in vivo* and they generate gliomas with a high penetrance (see Materials and Methods). As expected, overexpression of Tau clearly inhibited the growth of SVZ-EGFRamp/wt mouse gliomas (Fig. 4I) but it had no effect on SVZ-EGFRvIII tumors (Fig. 4J). Similar results were obtained when we overexpressed Tau in two human primary cell lines, RG1 (*EGFR*amp) and 12o150 (*EGFR* amplified and mutated, *EGFR*mut). Tau expression impaired the growth of the *EGFR*amp cell line (Fig. 4K) but it had no effect on *EGFR*mut cells (Fig. 4L). Remarkably, Tau did not inhibit the survival or the self-renewal capacity of the mouse (fig. S5E and F) or the human (fig. S5G and H) glioma cells *in vitro*, suggesting that the consequence of Tau overexpression on EGFR signaling pathway must be only relevant in the context of the tumor microenvironment (TME).

### Microtubule stabilizators like Tau and EpoD favor the degradation of wild-type EGFR

When we analyzed the dissected tumors after Tau overexpression we observed that the levels of phospho-EGFR were attenuated in SVZ-EGFRamp/wt tumors (Fig. 5A) as well as in *EGFR*amp xenografts (Fig. 5B), whereas there was no change in SVZ-EGFRvIII gliomas (Fig. 5C) or in *EGFR*mut xenografts (Fig. 5D). These data reinforce the notion that Tau opposes wild-type but not mutant EGFR signaling.

**Fig. 5.**
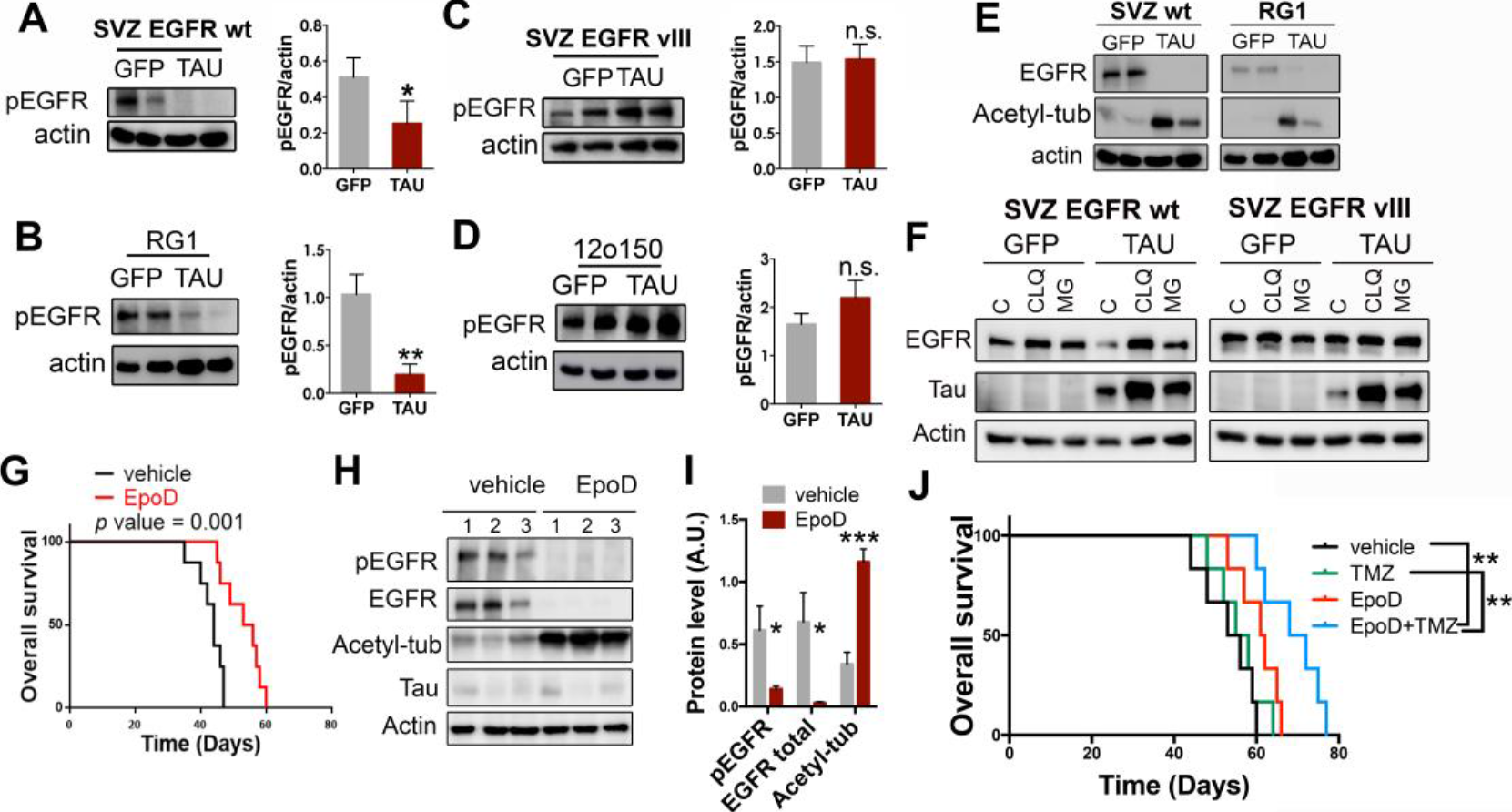
Tau and microtubule-stabilizing compounds impair wild-type EGFR stability and signaling in gliomas. (**A** to **D**) WB analysis and quantification of phosphorylated EGFR (pEGFR) in SVZ EGFRwt/amp (A), RG1 (B), SVZ EGFR vIII (C) and 12o150 (D) tumors after the overexpression of GFP or Tau. Actin was used as a loading control. (**E**) WB analysis of total EGFR and Acetylated tubulin (Acetyl-tub) in SVZ EGFR wt/amp and RG1 tumors after the overexpression of GFP or Tau. Actin was used as a loading control. (**F**) WB analysis of total EGFR in SVZ EGFRwt/amp and SVZ EGFR vIII cells after the overexpression of GFP or Tau, in the absence or in the presence of MG123 (MG, 10 μM) and Chloroquine (CLQ, 200 μM). (**G**) Kaplan-Meier overall survival curves of mice that were orthotopically injected with RG1 cells and subsequently treated with intraperitoneal injections (2/week) of EpoD (1 mg/Kg) (n=8). (**H** and **I**) WB analysis and quantification of pEGFR, total EGFR and Acetyl-tub in the tumors (G). Actin was used as loading control. (**J**) Kaplan-Meier overall survival curves of mice that were orthotopically injected with RG1 cells and subsequently treated with intraperitoneal injections (2/week) of EpoD (1mg/Kg) and/or TMZ (5mg/Kg/day) (n=6). *, *p* ≤0.05, **, *p* ≤0.01, ***, *p* ≤0.001. n.s. non significant.

Previous results from our group have demonstrated that, as a result of Tau function, stabilized microtubules become heavily acetylated through the inhibition of HDAC6 (histone deacetylase 6). In fact, the increase in tubulin acetylation can serve as a readout of Tau expression (16). Moreover, it has been proposed that this acetylation promotes the subsequent degradation of EGFR due to changes in the microtubule-dependent endocytic machinery (17). In agreement with these published data, we observed high levels of acetylated tubulin in Tau overexpressing gliomas (mouse or human), which were paralled by a strong decrease in the amount of total EGFR protein (Fig. 5E). Interestingly, the downregulation of EGFR protein can be observed *in vitro* after Tau overexpression, and this effect was reverted in the presence of MG132 (proteasome inhibitor) or chloroquine (lysosome inhibitor) (Fig. 5F). Collectively, these results suggest that Tau impairs EGFR signaling through the stabilization of the microtubules and the subsequent alteration of the receptor trafficking, which leads to its degradation. To validate this hypothesis we treated RG1(*EGFR*amp)-injected mice with a Taxol derivative, Epothilone D (EpoD). This kind of components can stabilize the microtubules as well. In fact, they compete with Tau for the same tubulin-binding site. EpoD in particular has been proved to reach the brain and revert some of the axonal defects of a Tau loss-of-function model (18). Systemic treatment with EpoD significantly delayed RG1 tumor formation (Fig. 5G) and reduced the amount of phospho-EGFR in the tumors (Fig. 5H and I) without changing the levels of Tau protein (Fig. 5H). This was accompanied by a strong downregulation of the levels of EGFR and a clear increase in the amount of acetylated tubulin (Fig. 5H and I). Altogether, our data support the notion that the microtubule-related function of Tau is responsible for its effect on EGFR signaling and tumor growth in gliomas. Moreover, they suggest that taxol derivatives could reduce the aggressiviness of gliomas. In fact, we observed that EpoD treatment made RG1 tumors more sensitive to chemotherapy (Fig. 5J) so there could be a therapeutic opportunity for these compounds to be used in glioma patients.

### Tau blocks the mesenchymalization of EGFRamp/wt glioma cells

The bioinformatic analysis showed that the levels of *Tau (MAPT)* correlate positively with the PN signature while they exhibit a strong inverse correlation with the MES expression profile of gliomas (Fig. 6A, and fig. S6A to F). In addition, we observed that *Tau* expression inversely correlates with the NF-κB (Nuclear factor kappa-light-chain-enhancer of activated B cells) and the inflammatory pathways, but not with other signatures, like those associated with hypoxia or PDGFRα (Fig. 6A and B). These in silico observations were confirmed by the WB analysis of RG1 (EGFRamp) xenografts, which showed that overexpression of Tau induces a clear inhibition of the NF-κB subunit phosphorylation, as well as a strong reduction in the protein level of TAZ, a master regulator of the MES phenotype (19) (Fig. 6C and D). Similar results were obtained in the WB analysis of 12o15 xenografts (Fig. 6E and F), a primary cell line that overexpresses the receptor in the absence of gene amplification (20). However, Tau did not change the expression of other MES genes (Fig. 6G) and it had no effect on TAZ expression or NF-κB activation in *EGFR*mut tumors, where there was no inhibition of the receptor upon Tau induction (fig. S6G). These data suggest that EGFR signaling induces the MES features in gliomas, which can be impaired by Tau, provided that no additional mutations in the receptor are present.

**Fig. 6.**
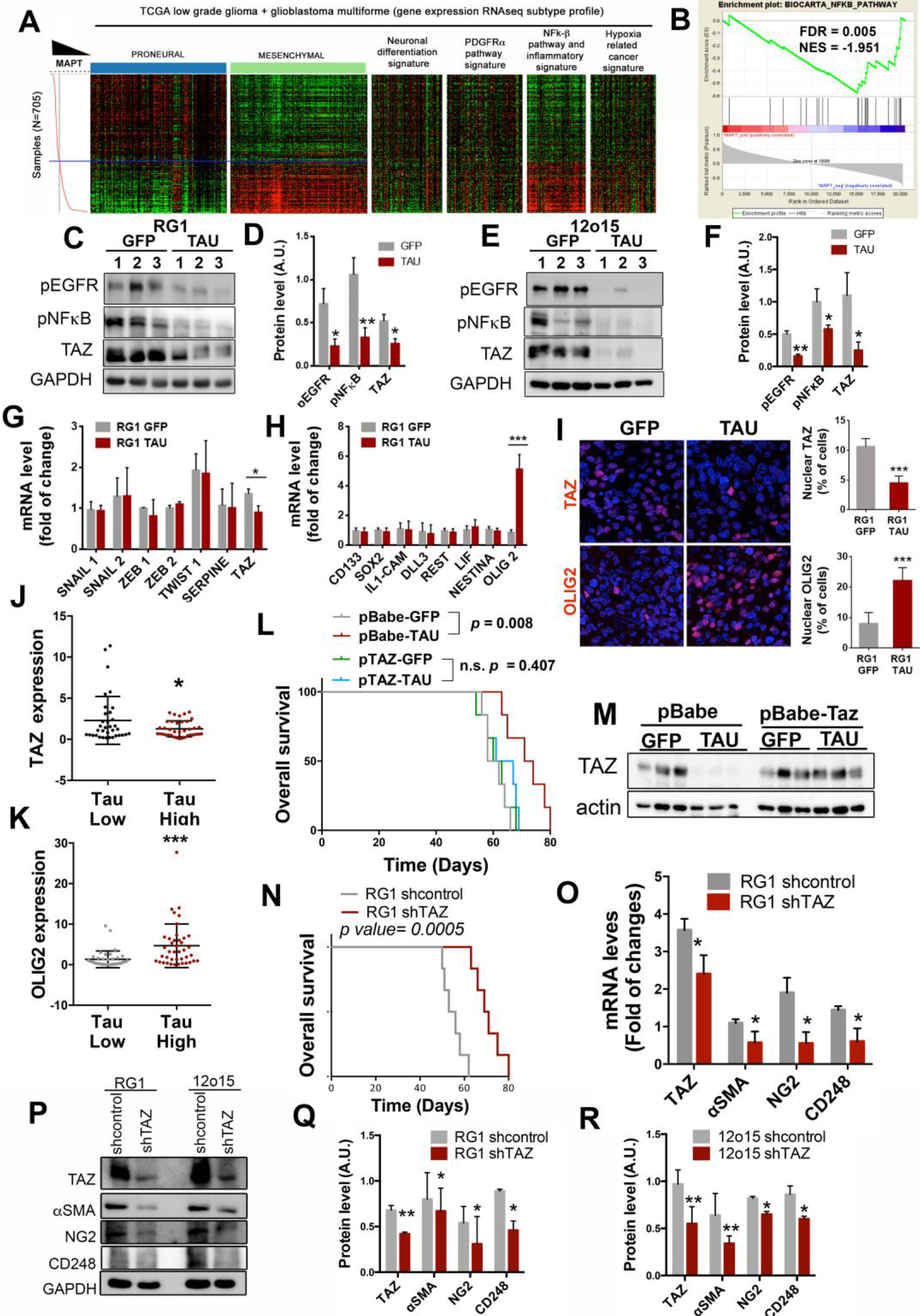
Tau blocks the mesenchymal features of *EGFR* amp/wt cells through the inhibition of the EGFR/NF-κB/TAZ axis. (**A**) Heatmap of PN, MES, neuronal differentiation, PDGFRα pathway, NF-κB/ inflammation, and hypoxia-related cancer expression genes depending on *Tau (MAPT)* expression levels. (**B**) GSEA enrichment plot analysis using Tau gene expression values as template and the NF-κB pathway geneset from the Biocarta pathways database. (**C** to **F**) WB analysis (**C** and **E**) and quantification (**D** and **F**) of phosphorylated EGFR (pEGFR), phosphorylated pNF-κB (p65) and TAZ in RG1 (**C** and **D**) and 12o15 (**E** and **F**) xenografts expressing either GFP or Tau. GADPH was used as a loading control. (**G** and **H**) qRT-PCR analysis of MES (G) and PN-subtype (H) related genes in RG1 xenografts expressing GFP or Tau. *HPRT* was used for normalization. (**I**) Representative images of TAZ (top) or OLIG2 (bottom) IF staining of sections from GFP or Tau overexpressing RG1 gliomas. Quantification is shown on the right. **(J** and **K**) qRT-PCR analysis of *TAZ* (J) and *OLIG2* (K) in gliomas. Tumors were classified in two groups based on the expression of *Tau (MAPT)*. (n=72). *HPRT* expression was used for normalization. (**L**) Kaplan-Meier overall survival curves of mice that were orthotopically injected with RG1 cells overexpressing GFP, Tau or GFP, Tau plus TAZ (n=6). (**M**) WB analysis of TAZ in the tumors (L). Actin was using as loading control. (**N**) Kaplan-Meier overall survival curves of mice that were orthotopically injected with RG1 shControl or RG1 shTAZ cells (n=6). (**O**) qRT-PCR analysis of pericytic–related genes in RG1 shControl and RG1 shTAZ xenografts. (**P** to **R**) WB analysis (**P**) and quantification (**Q** and **R**) of TAZ, αSMA, NG2 and CD248 in RG1 (P and Q) or 12O15 (P and R) cells after TAZ downregulation. GADPH levels were used for normalization. *, *p* ≤0.05; ***, *p* ≤0.001.

Tau overexpression also induced the expression of *OLIG2* (Fig. 6H), a transcription factor that is frequently used as a *bona fide* PN marker (21). Moreover, the IF analysis of RG1 gliomas confirmed that Tau represses the nuclear expression of TAZ, whereas it induces the nuclear expression of OLIG2 in the tumor cells (Fig. 6I). These data were confirmed by RT-PCR analysis of human samples (Fig. 6J and K), and by in silico studies (fig. S6H), which showed that Tau correlates inversely with *TAZ* and positively with *OLIG2* expression. Altogether, our results indicate that Tau induces a change in the glioma phenotype, repressing MES regulators and inducing PN promoters in the tumor cells through the regulation of the EGFR/TAZ/NF-kB pathway. It is important to remakr that Tau did not change the tumor burden of RG1 cells in the presence of TAZ overexpression (Fig. 6L and M), which suggest that Tau represses the growth of EGFRamp/wt glioma growth by blocking the plasticity of tumor cells and the appearance of mesenchymal features.

Previous results from our group and others have shown that, in comparison with PN cancer stem cells (CSCs), MES CSCs are more capable of glioma initiation, probably associated with their increased angiogenic capacity (22,23). In agreement with this observation, the overexpression of Tau in RG1 cells impaired their capacity to form subcutaneous xenografts (fig. S7A and B). Furthermore, we found that most of the genes upregulated with TAZ in gliomas were associated with angiogenesis and tumor vasculature (fig. S7C to F). Particularly, we noticed that markers of pericytes, which are considered as perivascular MES stem cells (24), were among the genes which correlated better with TAZ in gliomas (fig. S7G and H). In order to examine if this transcription factor could modulate the function and/or the number of mural cells we induced the expression of shTAZ in RG1 cells and we performed an ortothopic *in vivo* experiment. As expected, the knock down of this gene clearly reduced the aggressiviness of the tumors (Fig. 6N). Moreover, we observed a significant inhibition of the expression of several pericyte markers in RG1 shTAZ compared to control tumor samples (Fig. 6O). It is worth mentioning that TAZ seems to regulate peryctic differentiation in a cell-autonomous way, as we also observed a reduction of these markers after the *in vitro* expression of shTAZ in RG1 and 12o15 cells (Fig. 6P to R). These results are in accordance with some recent publications showing that most of the pericytic functions in GBM are performed by the highly plastic CSCs, which undergo a transdifferentiation process, acquiring mesenchymal and mural cell features (25–27).

### Tau reduces the amount of tumor-derived-pericytes and normalizes the vasculature of EGFRamp gliomas

Our results indicate that Tau impairs the mesenchymalization of glioma cells by blocking the expression of TAZ, which is induced in response to EGFR activation. Moreover, we could hypothesize that, as a result of this inhibition, there is an important blockade of the capacity of the tumor cells to transdifferentiate into pericytes. Therefore, we decided to analyze the vascular component of the tumors after the overexpression of Tau in RG1 (EGFRamp) cells. We observed a marked decrease in the number of αSMA and CD248 positive cells (Fig. 7A), together with a significant inhibition of the expression of human pericytic markers, in Tau-expressing tumors (Fig. 7B). A similar result was observed in 12o15 (EGFRwt) (fig. S8A), but not in 12o150 (EGFRmut) xenografts (fig. S8B). Remarkably, Tau did not impaired the expression of mouse pericityc genes (Fig. 7C), reinforcing the idea that this protein inhibits the glioma-to-pericyte transdifferentiation but it does not influence the amount and/or the differentiation capacity of host pericytes.

**Fig. 7.**
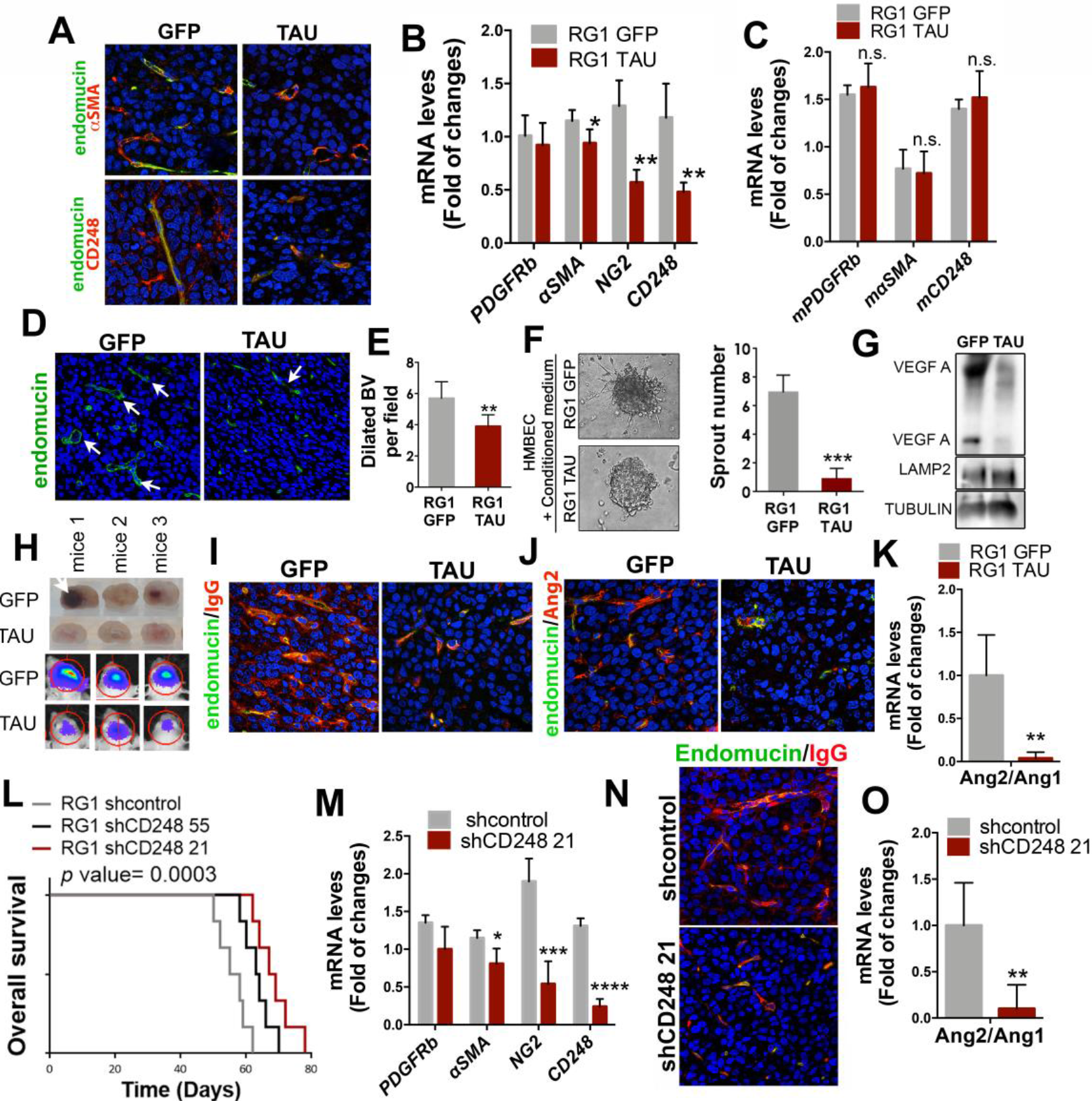
Tau overexpression blocks the appearance of tumor-derived-pericytes in *EGFRwt/*amp gliomas, inhibiting angiogenesis and normalizing the tumor vasculature. (**A**) Representative images of endomucin and αSMA (top) or endomucin and CD248 (bottom) IF co-staining of sections from RG1 xenografts expressing GFP or Tau. (**B**) qRTPCR analysis of pericytic–related genes (using human-specific primers) in RG1 GFP and Tau xenografts. *HPRT* was used for normalization. (**C**) qRT-PCR analysis of pericytic–related genes (using mouse-specific primers) in RG1 GFP and Tau xenografts. *Actin* was used for normalization. (**D**) Representative images of endomucin IF staining of sections from RG1 xenografts expressing GFP or Tau. (**E**) Quantification of dilated blood vessels in (D) (indicated by arrows). (**F**) Representative contrast phase images of HMBEC cells cultured in the presence of conditioned media from RG1 GFP or RG1 Tau cells. The number of sprouts is shown on the right. (**G**) WB of VEGF in the supernatant of RG1 cells after overexpression of GFP or Tau. LAMP2 and TUBULIN were used as loading controls. (**H**) Representative images of sectioned mouse brains (top) and pseudo-color images of Fluc bioluminescence (bottom), after the injection of GFP or Tau-expressing RG1 cells. (**I** and **J**) Representative images of IF co-staining of endomucin and IgG (I) or endomucin and Ang2 (J) on RG1 xenografts expressing GFP or Tau. (**K**) Ratio of *Ang2/Ang1* expression levels measured by qRT-PCR in RG1 gliomas after Tau overexpression. *Actin* was used for normalization. (**L**) Kaplan-Meier overall survival curves of mice that were orthotopically injected with RG1 cells expressing shcontrol, shCD248 55 or shCD248 21 (n=6). (**M**) qRT-PCR analysis of pericytic-related genes in RG1 tumors expressing shcontrol or shCD248 21. (**N**) Representative images of IF co-staining of endomucin and IgG from RG1 xenografts expressing shControl or shCD248 21. (**O**) Ratio of *Ang2/Ang1* expression levels measured by qRT-PCR in RG1 gliomas after *CD248* downregulation. *Actin* was used for normalization. *, *p* ≤0.05; **, *p* ≤0.01; ***, *p* ≤0.001; ****, *p* ≤0.0001. n.s., nonsignificant.

The IF analysis of the tissues also revealed that there is an important inhibition of cellular proliferation (fig. S8C and D) after Tau overexpression, accompanied by a significant decrease in the number of dilated blood vessels (Fig. 7D and E), typical of malignant gliomas (28). The angiogenic signals that are reduced upon Tau induction must be derived, at least in part, from the tumor cells, as the supernatant of RG1 cells overexpressing Tau has a reduced capacity to induce HMBEC (Human Brain Microvascular Endothelial Cells) sprouting (Fig. 7F) and contains less VEGF (Vascular Endothelial Growth Factor) (Fig. 7G).

We had previously noticed the lack of hemorrhages in *EGFR*amp tumors overexpressing Tau, an event that was frequent in control tumors (Fig. 7H). In parallel, we observed a reduction in the extravasation of IgG after Tau overexpression (Fig. 7I). These observations suggest that tumors that overexpress Tau present a less aberrant tumor vasculature, with an increase in the intregrity of the blood brain barrier (BBB). In agreement with this hypothesis, Tau induced a clear reduction in the levels of Ang2 (Angiopoietin 2) (Fig. 7J), a key regulator of angiogenesis that has been associated with vascular abnormalities in glioma, acting as an antagonist of Ang1 (29). In fact, we observed a significant decrease in the *Ang2/Ang1* ratio after Tau overexpression (Fig. 7K) in the *EGFR*amp model, but not in the *EGFR*mut xenografts (fig. S8E). These results suggest that the transdifferentiation of glioma cells into mesenchymal/pericyte-like cells, induced by EGFR signaling, favors the secretion of angiogenic signals and the formation of an aberrant vasculature. Interestingly, this process can be impaired by Tau in EGFRwt/amp, but not in EGFRmut gliomas.

In order to confirm the relevance of the tumor-derived-pericytes in the growth of EGFRamp gliomas, we downregulated *CD248* expression in RG1 cells. CD248, also called endosialin, aids in supporting tumor microvasculature and it is expressed in pericytes, especially in malignant solid tumors, including high-grade gliomas (26,30). We picked the most effective shRNA sequences (fig. S8F) and we injected the interfered cells into the brains of immunodeficient mice, in the presence of normal host pericytes. We observed a significant delay in tumor growth after *CD248* downregulation (Fig. 7L).

Moreover, the inhibition of *CD248* in the tumor cells reduced the expression of other pericytic markers *in vitro* (fig. S8G) and *in vivo* (Fig. 7M). These results suggest that CD248 is a key player in the transdifferentiation of tumor cells into pericytes. On top of that, tumors formed after *CD248* downregulation showed a reduction in the extravasation of IgG (Fig. 7N) as well as a decreased *Ang2/Ang1* ratio (Fig. 7O), wich further supports the hypothesis that the depletion of tumor-derived-pericytes (at least in EGFRwt/amp tumors) normalizes the glioma vasculature and reduces the aggressiviness of this type of cancer.

### Tau expression is a surrogate marker of the less aggressive vascular behavior of gliomas

The data presented so far place Tau at the boundary between the most common genetic alterations of gliomas (those that affect *IDH1/2* and *EGFR*) and the regulation of the tumor microenvironment, in particular the vascular phenotype. In order to translate the results obtained in mouse models into the clinical settings, we used our own cohort of patient’s samples (RT-PCR analysis) and the data from the TCGA (in silico analysis). We divided the tumors into an IDH1 mut (those with higher transcript levels of Tau) and an IDH1 wt group, and we subclasified the second one into High or Low Tau gliomas (Fig. 8A and B). First, we confirmed that the inverse correlation between the levels of *Tau* and the amount of phospho-EGFR was maintained in the human samples (Fig. 8C). Moreover, we also observed a significant inverse correlation between the transcript levels of *CD248* and the expression of *Tau* (Fig. 8D and E), which was confirmed by the IHC analysis of the tumors. The representative images in Fig. 8F evidence the gradual normalization of the vasculature in parallel with the increase in Tau levels (Fig. 8F). Furthermore, the quantification of the CD248 score (Fig. 8G), the amount of *CD34* (Fig. 8H) and the amount of dilated vessels (Fig. 8I), confirmed the visual differences and reinforced our model, in which Tau expression could function as a surrogate marker of the less aggressive vascular behavior in gliomas.

**Fig. 8.**
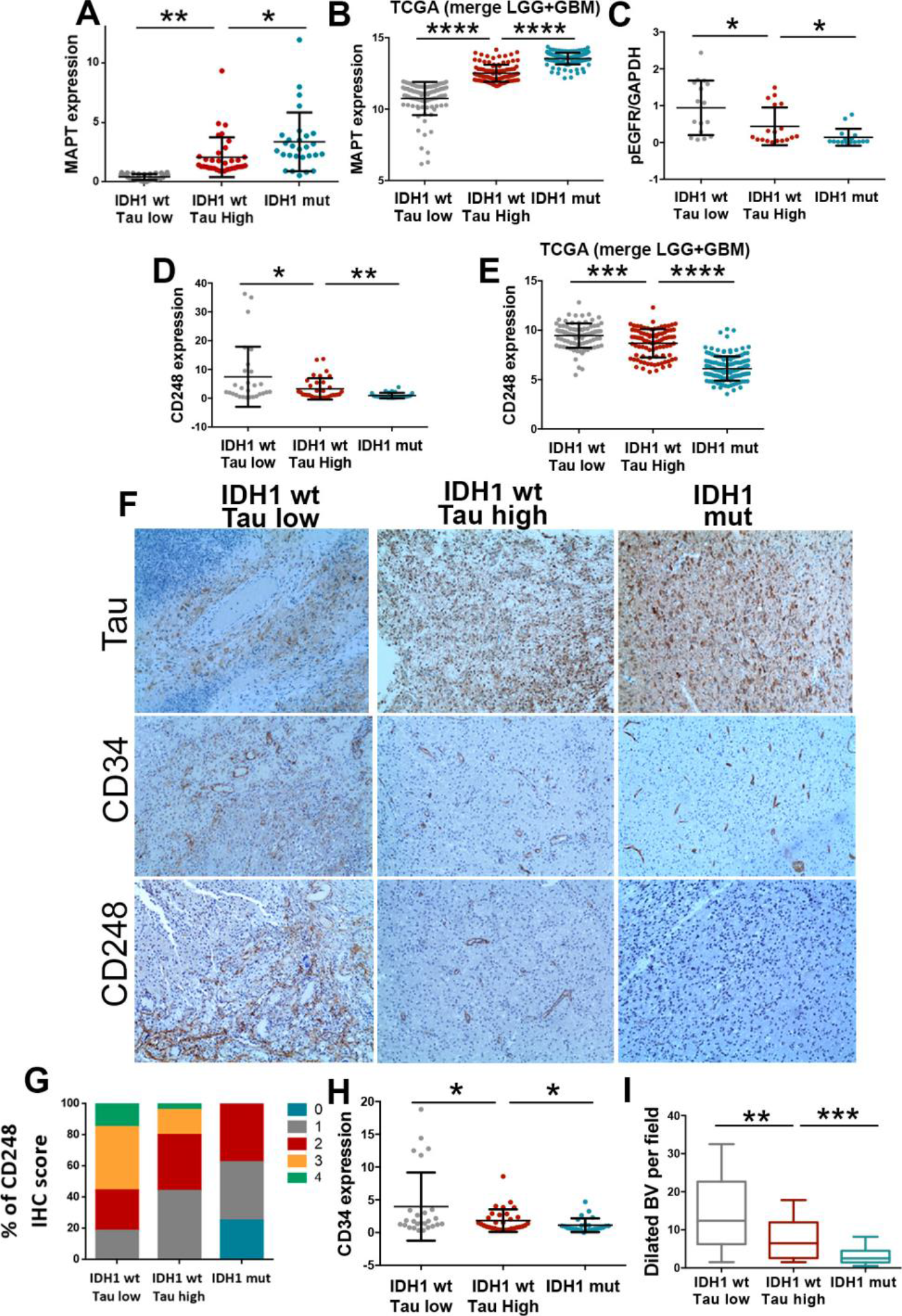
Analysis of Tau and vascular molecules in human samples. (**A** to **E**) *Tau (MAPT)* (A and B), and pEGFR (C), and CD248 (D and E) expression levels were determined by qRT-PCR analysis (Glioma cohort) (n=87) (A to C) or RNAseq analysis (TCGA-LGG+GBM cohort) (n=319) (B and E). Tumors were classified in three groups: IDH1 wt (Tau Low), IDH1 wt (Tau High) and IDH1 mut. *HPRT* expression was used for normalization. (**F**) Representative pictures of the IHC staining of Tau, CD34 and CD48 in three tumors, one for each of the groups. (**G**) Percentage of tumors with different CD248 IHC score in E (n=68). (**H**) CD34 levels were determined by qRT-PCR analysis (GBM cohort) (n=87). (**I**) Quantification of the dilated blood vessels in E (n=68). *, *p* ≤0.05; **, *p* ≤0.01; ***, *p* ≤0.001; ****, *p* ≤0.0001. n.s., non-significant.

## DISCUSSION

IDH1/2 mutations identify a genetically and clinically distinct glioma entity. Certainly, patients with such tumors have a much better prognosis, independently of the histological classification, and they show improved responses to chemotherapy and/or irradiation (31,32). Indeed, the re-expression of IDH1 R132H in GBM can reduce tumor growth (fig. S4F) (13,33). However, the molecular basis for the tumor-suppressor functions of IDH mutations are unclear. Here, we have identified Tau, a known microtubule stabilizer, as a new epigenetic target of IDHmut in gliomas. Based on our results we can propose that these mutant enzymes, acting through the increase in Tau expression, favor the normalization of the vasculature and impede the progression of the disease (fig. S9). In fact, Tau downregulation induces a dramatic increase in tumor burden in the orthotopic xenografts. Moreover, the longitudinal analysis of a set of paired samples showed that Tau expression decreases as LGG evolve into higher grade lesions. Interestingly, IDH1/2 status seems to be consistent during glioma progression (34,35), suggesting that some other mechanism is inhibiting Tau transcription in the recurrent tumors. One possible explanation would be that an increase in the levels of wild-type IDH protein in the recidives could impair Tau expression, as it happens in the mouse xenografts (Fig. 4D). Indeed, it has been recently shown that the non-mutated IDH1 is upregulated in primary GBM in comparison with secondary GBM or LGG, where it promotes aggressive growth and therapy resistance (5).

From the moment that IDH1/2 mutations were identified in gliomas, it seemed clear that they represent early events in the process of tumorigenesis (36). A decade later we know that these proteins induce important metabolic and epigenomic changes, which could explain their oncogenic properties (4,13,14,37,38). Consequently, a plethora of inhibitors have been developed to block IDHmut function in different cancers. Indeed, some of them have shown anti-tumor acitivty in mouse glioma models (39) and they have been approved to be tested in IDHmut patients (40). However, the tumor-suppressor functions of these proteins in gliomas, including the vascular normalization that we have described here, could represent a severe pitfall for these approaches. In fact, the treatment with IDHmut inhibitors has been associated with a radioprotective effect (41) and a decreased sensitivity to cisplatin (42). Regarding IDHwt gliomas, our results suggest that microtubule stabilizers could imitate Tau function and reduce the aggressiveness of the tumors, which could render them sensitive to conventional chemotherapy (Fig. 5I). The same compounds could be tested in combination with IDHmut inhibitors in LGG, in an attempt to impair tumor growth and, at the same time, block the transformation into a more aggressive tumors.

Our results indicate that Tau is a key inhibitor of wild-type EGFR signaling in gliomas. Accordingly, drugs that interfere with microtubule function have been associated with EGFR inactivation in other cancers (43), as they seem to alter endocytic trafficking (11). Moreover, it has been shown that microtubule acetylation, which is promoted by Tau function (16), promotes the degradation of EGFR through changes in the microtubule-dependent endocytic trafficking (17). We propose that a similar mechanism could be operating in gliomas, but only in the absence of EGFR mutations. In that sense, there is plenty of literature that suggest that EGFR mutant proteins, especially the vIII isoform, can maintain the signal in the absence of the ligand (44), possibly due to changes in the turnover of the mutant receptor (45). Moreover, it has been proposed that some of the EGFR downstream targets, like NF-κB, become constitutively activated in GBM after vIII expression (46). The data presented here further supports that EGFR mutations induce a constitutive activation of NF-κB, which would be unsensitive to microtubule modulators.

Downstream of EGFR/NF-κB signaling, we have found that Tau overexpression severely reduced the amount of TAZ in GBM. TAZ is a transcriptional coactivator that play a prominent role in gliomas (47), controlling the MES signature (19). Moreover, TAZ is induced by NF-κB activation in gliomas and promotes radio-resistance (19). It has been recently proposed that *TAZ* promoter is a direct target of NF-κB (48) so Tau could be inhibiting indirectly the transcription of the *TAZ* gene. However, we cannot discard a possible regulation of the protein stability downstream of EGFR/NF-κB signaling.

It is well known that IDH mutant gliomas are characterized by a lesser extent of contrast enhancement than wild-type tumors, even if we only consider GBM (49). However, the molecular explanation for this behavior was still missing. Our results show that Tau, acting downstream of IDHmut, could normalize the blood vessels and reduce the BBB leakage. These changes could cooperate or even precede other alterations observed in the TME of IDHmut gliomas, like the absence of microthrombi (50) as well as the decrease in necrosis areas (49) and hypoxia-induced angiogenesis (51). Moreover, the immune component and its pro- or anti-tumoral properties could also be affected by the vascular phenotype of IDHmut gliomas (52), explaining the decrease in the immune infiltrate observed in mouse models and human samples (33). Future experiments are warranted in order to decipher the participation of Tau in these other phenotypes.

Although cancer and neurodegenerative diseases are considered as opposite phenomena, there could be a positive association between AD prevalence and GBM incidence (12). In AD and other tauopathies, there is an accumulation of Tau aggregates (gain of toxic function). However, this aggregation of Tau compromises its microtubule-stabilising functions (loss of physiological function), favoring the evolution of the pathology. In agreement with the loss-of-function model, we have reported behavioral changes and neurogenesis in aged Tau knockout mice (53). Interestingly, EMT genes are enriched in the affected areas of AD brains (54). Moreover, there is an important neurovascular dysfunction at early stages of AD, associated with BBB breakdown and inflammation (55), which could be linked to pericyte loss (56). Although there is still missing evidence of the relevance of Tau (especially the astrocytic Tau) in these phenotypes, it is tempting to speculate that the loss of function of this protein could represent a link between the progression of the two types of brain diseases. If this hypothesis is correct, targeting pericytes or the mesenchymal transdifferentiation of the astrocytes could be an interesting therapeutic approach to be tested in several neurological disorders. Moreover, it would be worth studying if AD related drugs could be repurposed to treat brain tumors. In that sense, we have demonstrated that EpoD, a microtubule stabilizing agent that reduces AD pathology in mouse models (18), slows down *in vivo* glioma growth without any apparent secondary effect. Another second-generation taxane, cabazitaxel, has demonstrated a good anti-glioma activity in preclinical tests (57) and is being evaluated as a novel cytotoxic therapy (NCT01740570, NCT01866449). However, we propose that lower doses of these compounds could imitate the effect of Tau overexpression, reducing the aggressiviness of gliomas with less secondary effects for the patients. By contrast, as they compete for the same microtubule binding site of Tau, high levels of this protein could identify gliomas that would be resistant to taxol derivatives, as it has been shown for other cancers (58).

Collectively, our results provide and explanation for the better outcome of IDHmut gliomas, which could have important implications for several aspects of glioma research and clinical practice. Moreover, the understanding of how Tau governs the vascular component of the TME in gliomas could provide novel explanations for the neurovascular dysfunction observed in AD and other dementias.

## MATERIALS AND METHODS

### Human samples

Glioma tissues embedded in paraffin were obtained after patient’s written consent and with the approval of the Ethical Committees of “Hospital 12 de Octubre” (CEI 14/023) and “Hospitales de Madrid” (14.10.632-GHM, 18.12.1337-GHM) (Table S1 and 2).

### Human and mouse glioma cells

RG1 cells were kindly donated by Rosella Galli (San Raffaele Scientific Institute). The rest of the human cells (Table S3) were obtained by dissociation of surgical specimens from patients treated at the “Hospital 12 de Octubre” (Madrid, Spain). They were obtained after patient’s written consent and with the approval of the Ethical Committee (CEI 14/023). They belong to the Biobank of that Hospital. Cells were grown in Complete Media (CM): Neurobasal supplemented with B27 (1:50) and GlutaMAX (1:100) (Thermo-Fisher-Scientific); penicillin-streptomycin (1:100) (Lonza); 0.4% heparin (Sigma-Aldrich); and 40 ng/ml EGF and 20 ng/ml bFGF2 (Peprotech). Mouse SVZ models were obtained by retroviral expression of EGFRwt or EGFRvIII in primary neural stem cell (NSC) cultures from p16/p19 ko mice. The NSCs were obtained as previously described (59) and they were grown in CM. After infection, the cells were injected into nude mice and the tumors that grew were dissociated and established as SVZ-EGFRamp/wt or SVZ-EGFRvIII models. Both models express GFP and luciferase as reporters. These established cell lines (300,000 cells) give rise to gliomas when they are implanted in the brains of nude mice with a 100% penetrance. The average survival is around 62 days for the SVZ EGFR wt/amp model and around 32 days for the SVZ EGFR vIII cells, showing a proliferation rate of more than 10% in SVZ EGFR wt/amp tumors and more than 30 % for SVZ EGFR vIII tumors. Both mouse glioma models, SVZ EGFR wt/amp and SVZ EGFR vIII, showed a complete dependence on EGFR activity, since their growth can be impaired by a tyrosin kinase inhibitor (dacomitinib) (44). NPA (NRAS, shP53, shATRX) IDH1 wt and NPA (NRAS, shP53, shATRX) IDH1 R132H were provided by Maria G. Castro (University of Michigan) (13) and cultured in CM.

### DNA constructs and lentiviral/retroviral production

The lentivirus that encodes the longest wild type isoform of Tau in human brain harboring four repeats and two N-terminal inserts followed by GFP linked by an IRES was a gift from by Prof. Kenneth S. Kosik (UC Santa Barbara). As a control we used a lentivector encoding E-GFP, pRRLSIN.cPPT.PGK-GFP.WPRE (Addgene plasmid 12252). pLV-Hygro-Luciferase (VectorBuilder #VB150916-10098) was used as reporter. Lentiviral vector to express shRNAs were: shCD248 (Sigma #SHCLNG-NM_020404: TRCN0000053455, TRCN0000053457, TRCN00000443679, TRCN00000429396, TRCN0000043782) and shTAZ (TRCN0000370006, TRCN0000370007). Retroviral vectors used were pBabe-EGFR wt (#11011), MSCV-XZ066-GFP-EGFR vIII (#20737), pBabe-puro-Flag-IDH1 (#62923), pBabe-puro-Flag-IDH1-R132H (#62924) (Addgene) and pBabePuroTAZ-WT was a generous gift from Kun-Liang Guan. To obtain the virus, the 293T cells were transiently co-transfected with 5 μg of appropriate lentivector plasmid, 5 μg packaging plasmid pCMVdR8.74 (Addgene #Plasmid 22036) and 2 μg VSV-G envelope protein plasmid pMD2G (Addgene #Plasmid 12259) using Lipofectamine Plus reagent (Invitrogen). Retrovirus and lentivirus supernatant was prepared by transfection of 293T cells and collection of the supernatant 48 hr after.

### Intracraneal tumor formation and treatment in vivo

Animal experiments were reviewed and approved by the Research Ethics and Animal Welfare Committee at “Instituto de Salud Carlos III” (PROEX 244/14 and 02/16), in agreement with the European Union and national directives. Intracranial transplantation to establish orthotopic xeno- and allo-grafts was performed injecting 100.000-300.000 cells (resuspended in 2 μl of culture stem cell medium) with a Hamilton syringe into athymic Nude-Foxn1nu brains (Harlan Iberica). The injections were made into the striatum (coordinates: A–P, −0.5 mm; M–L, +2 mm, D–V, −3 mm; related to Bregma) using a Stoelting Stereotaxic device. When applicable, tumor growth was monitored in an IVIS equipment (Perkin Elmer) after intraperitoneal injection of D-luciferin (75 mg/Kg) (PerkinElmer). The animals were sacrificed at the onset of symptoms. Mice were treated with Epothilone D (Abcam, ab143616) (1 mg/kg two days per week through intra-peritoneal injection) and/or Temozolomide (Sigma Aldrich) (5mg/kg daily through intraperitoneal injection). Epothilone D was dissolved in 4% DMSO +10%Polysorbate. Temozolomide was dissolved in 1% BSA+PBS. Control animals were treated with these solvents.

### Growth curve and sphere formation assay

Glioma cells were infected by the control lentivirus (LV-GFP) or lentivirus directing expression of TAU (LV-TAU-GFP). The tumor spheres were Accumax-dissociated to single cells, and 500/1000 cells of each condition were plated in a p24-well-plate in triplicate. Five days after plating, spheres and cell number were measured.

### EGFR degradation experiment

GFP or TAU SVZ EGFRwt/amp cells were grown in stavirng media for 1h and then EGF (100 ng/ml) was added and the cells were incubated in the presence of DMSO, MG132 or Chloroquine for 2h. Then, cells were collected and lysed and subsequently analyzed by western blot, as described below.

### Endothelial cell sprouting assay

In order to generate endothelial cell spheroids, 1×10^6^ cells of HMBEC were suspended in CM and seeded in nonadherent plates. These spheroids were harvest within 48 hours and embedded into matrigel in 96-well plates with conditioned medium generated for each cell line. After 24 h, sprouting was induced and the number of sprouts for each spheorid were quantified. For the generation of the conditioned media, RG1 (GFP or Tau) were grown during 48 hours in serum-free culture media (DMEM-F12 supplemented with FGF2 (50ng/ml) and penicillin-streptomycin). The medium was filtered with a 70-μm filter before use.

### In vivo limiting dilution assay

Increasin numbers of RG1 (GFP or TAU) cells were re-suspended in CM and Matrigel (BD) (1:10) and injected subcutaneously injected in nude mice. Animals were sacrificed before tumors reached a 1.5 cm in diameter. The statistical significances were calculated using the Extreme Limiting Dilution Analysis software (http://bioinf.wehi.edu.au/software/limdil/index.html).

### Inmunofluorescent (IF) and Inmunohistochemical (IHC) staining

Slides were heated at 60ºC for 1 hour followed by deparaffinization and hydration, washed with water, placed into antigen retrieval solution (pressure cooking) in 10 mM sodium citrate pH 6.0. Paraffin sections were permeabilized with 1% Triton X-100 (Sigma-Aldrich) in PBS and blocked for 1 hour in PBS with 5% BSA (Sigma), 10% FBS (Sigma) and 0,1% Triton X-100 (Sigma). The following primary antibodies (Table S4) were incubated O/N at 4°C. The second day, sections were washed with PBS three times prior to incubation with the appropriate secondary antibody (Table S4) (1:200 dilution) for 2h at room temperature. Prior to coverslip application, nuclei were counterstained with DAPI and imaging was done with Leica SP-5 confocal microscope. Otherwise, IHC sections were incubated with biotinylated secondary antibodies (1:200 dilution). Target proteins were detected with the ABC Kit and the DAB kit (Vector Laboratories).

### IHC quantification

The IHC score was judged from 0 (no staining) to 3 (Tau staining) or 4 (CD248 staining) on those samples with the strongest positive staining. For the longitudinal analysis of the primary and relapsed tumors we calculated the score of 10 high magnification pictures of each sample. The data depicted in Figure 2e,f is the average of these 10 pictures. For the quantification of the vasculature, we counted the number of dilated vessels per high-magnification field.

### Western Blot analysis

Protein content was quantified using BCA Protein Assay Kit (Thermo-Fisher-Scientific). Approximately 20 μg of proteins were resolved by 10% or 12% SDS-PAGE and they were then transferred to a nitro cellulose membrane (Hybond-ECL, Amersham Biosciences). The membranes were blocked for 1 h at room temperature in TBS-T (10 mM Tris-HCl [pH 7.5], 100 mM NaCl, and 0.1% Tween-20) with 5% skimmed milk, and then incubated overnight at 4ºC with the corresponding primary antibody diluted in TBS-T. The primary antibodies and the dilutions are shown in Table S4. After washing 3 times with TBS-T, the membranes were incubated for 2 h at room temperature with their corresponding secondary antibody (HRP-conjugated anti mouse or anti rabbit, DAKO) diluted in TBS-T. Proteins were visible by enhanced chemiluminiscence with ECL (Pierce) using the Amersham Imager 680.

### qRT-PCR assay

RNA was extracted from the tissue using RNA isolation Kit (Roche). Total RNA (1μg) was reverse transcribed with PrimeScript RT Reagent Kit (Takara). Quantitative real-time PCR was performed using the Light Cycler 1.5 (Roche) with the SYBR Premix Ex Taq (Takara). The primers used for each reaction are indicated in Table S5. Gene expression was quantified by the delta-delta Ct method.

### EGFR Sequencing of primary GBMs

To identify point EGFR mutations, cDNAs from the different cell lines were sequenced using the primers indicated in Table S6. The sequences were aligned and collated with the EGFR transcript (NM_005228.4) in the NIH GenBank database using the multiple sequence alignment tool Clustal Omega (www.ebi.ac.uk/Tools/msa/clustalo/) and the Sequencing Analysis Software v5.3.1 from Applied Biosystems. The identified mutations were analyzed at cBioPortal (www.cbioportal.org).

### Gene expression and survival analyses

*Tau (MAPT)* gene expression and follow-up overall survival data from human glioma tumors corresponding to TCGA Glioblastoma (GBM) and Brain lower grade Glioma (LGG) data sets were downloaded respectively from cBioPortal (http://www.cbioportal.org/) and TCGA databases (http://tcga-data.nci.nhi.gov/docs/publications/lgggbm_2015) using UCSC cancer browser. Kaplan-Meier survival curves were done within TCGA-GBM, TCGA-LGG, TCGA-GBM-LGG, Rembrandt, Gravendeel, Ducray, Freije and Nutt cohorts upon stratification based into low and high groups using expression values from *Tau (MAPT)* gene. Significance of differences in survival between groups was calculated using the log-rank test. These data were obtained from UCSC Xena-Browser (https://xenabrowser.net) and Gliovis (http://gliovis.bioinfo.cnio.es). Classification into classical, mesenchymal, neural and proneural subtypes was retrieved from the TCGA GBM data set (https://www.ncbi.nlm.nih.gov/pubmed/24120142) together with Tau/MAPT expression values. Differences in Tau expression between mesenchymal and other groups were calculated using Student’s t-test. In David gene onotology analysis we have used a cluster of 1000 genes co-expressed with Tau/MAPT. They were chosen using the highest values of the Spearman’s correlations. Correlation between gene expression values of MAPT versus other genes was done using Pearson analysis. Gene Set Enrichment Analysis (GSEA) was computed into the TCGA-GBMLGG cohort (RNAseq (IlluminaHiSeq)) using Tau gene expression as a continuous class label and genesets from the “CGP: chemical and genetic perturbations” and “CP:BIOCARTA: BioCarta gene sets” (n=) from the MSigDB genesets database.

### Analysis of methylation of the MAPT gene and CTCF Chip-seq binding

DNA-methylation analysis in human glioma within the MAPT locus was done using the TCGA data (Illumina Infinium HumanMethylation450 platform). Methylation beta values from 56 probes within MAPT locus were retrieved from the Xena browser (https://xenabrowser.net/) together with MAPT gene expression values. Pearson correlation values between each methylation probe and gene expression values, calculated for all samples, was calculated and represented together with the CpG islands located at the MAPT locus (CpG302, CpG26 and CpG21). Both TCGA LGG-GBM and LGG cohorts were individually analyzed. Correlation values were independently calculated for IDH1 mutant or wild-type samples in the TCGA LGG cohort. ChIP-seq analysis using anti-CTCF antibody was performed from profiling in IDH1 mutant and wild-type glioma patient specimens and culture models (GSE70991). Briefly tdf files from GEO repository from both IDH1 mutant and wild-type samples were downloaded and visualized using IGV browser. CTCF occupancy at the CpG islands located from MAPT loci was visualized.

### Statistical analysis

For bar graphs, the level of significance was determined by a two-tailed un-paired Student’s t-test. The difference between experimental groups was assessed by Paired t-Test and one-way ANOVA. For Kaplan-Meier survival curves, the level of significance was determined by the two-tailed log-rank test. For correlation analysis between each gene, expression data were tested by Pearson’s correlation coefficient and Spearman’s correlation coefficient. All analyses were performed with the GraphPad Prism 5 software. P values < 0.05 were considered significant (*p < 0.05; **p < 0.01; *** p< 0.001; **** p< 0.0001; n.s., non-significant). All quantitative data presented are the mean ± SEM from at least three simples or experiments per data point. Precise experimental details (number of animals or cells and experimental replicates) are provided in the Figure legends.

## SUPPLEMENTARY MATERIALS

Fig. S1. Association of Tau levels with the clinical pathology of diffuse gliomas.

Fig. S2. Patients derived Xenograft (PDXs) show a specific expression of Tau.

Fig. S3. Tau expression correlates with the presence of IDH mutations.

Fig. S4. The expression of Tau is associated with the IDH mutant methylation phenotype.

Fig. S5. Association of Tau function with the EGFR pathway in gliomas.

Fig. S6. Association of Tau expression with the GBM subtypes and the NF-kB-TAZ axis.

Fig. S7. Vascular phenotypes associated with TAZ in gliomas.

Fig. S8. Implication of the tumor-derived-pericytes on the glioma vasculature and growth.

Fig. S9. The vascular phenotype of gliomas is determined by the genetic status of EGFR and IDH and the expression of Tau.

Table S1. Human samples

Table S2. Paired human samples

Table S3. GBM cell lines

Table S4. Antibodies

Table S5. qRT-PCR primers

Table S6. Sequecing primers

## Acknowledgments

The authors would like to acknowledge Rosella Galli for donating RG1, Rafael Hortigüela and the Confocal service personell, for their technical support. The graphical abstract was created with images adapted from Servier Medical Art by Servier. Original images are licensed under a Creative Commons Attribution 3.0 Unported License.

## Author Contributions

Conceptualization: RG, BSC, JA and PSG; Investigation: RG, BSC, ARB, BH, FJN, DGP, JGG and RGE; Formal Analysis: RG and RGE; Resources: VGE, JMS, AHL, AAS, JS and MGC; Writing-Original Draft: RG, BSC, JA and PSG; Writing-Review & Editing: VGE, AAS, JS, JMS, AHL, MGC and RGE; Funding Acquisition: RG, JA and PSG; Supervision: JA and PSG.

## Reference List

1. Louis, D.N., Perry, A., Reifenberger, G., von, D.A., Figarella-Branger, D., Cavenee, W.K., Ohgaki, H., Wiestler, O.D., Kleihues, P., and Ellison, D.W. 2016. The 2016 World Health Organization Classification of Tumors of the Central Nervous System: a summary. Acta Neuropathol. 131:803–820.

2. Yan, H., Parsons, D.W., Jin, G., McLendon, R., Rasheed, B.A., Yuan, W., Kos, I., Batinic-Haberle, I., Jones, S., Riggins, G.J. et al 2009. IDH1 and IDH2 mutations in gliomas. N Engl J Med 360:765–773.

3. Turcan, S., Rohle, D., Goenka, A., Walsh, L.A., Fang, F., Yilmaz, E., Campos, C., Fabius, A.W., Lu, C., Ward, P.S. et al 2012. IDH1 mutation is sufficient to establish the glioma hypermethylator phenotype. Nature 483:479–483.

4. Lu, C., Ward, P.S., Kapoor, G.S., Rohle, D., Turcan, S., Abdel-Wahab, O., Edwards, C.R., Khanin, R., Figueroa, M.E., Melnick, A. et al 2012. IDH mutation impairs histone demethylation and results in a block to cell differentiation. Nature 483:474–478.

5. Calvert, A.E., Chalastanis, A., Wu, Y., Hurley, L.A., Kouri, F.M., Bi, Y., Kachman, M., May, J.L., Bartom, E., Hua, Y. et al 2017. Cancer-Associated IDH1 Promotes Growth and Resistance to Targeted Therapies in the Absence of Mutation. Cell Rep. 19:1858–1873.

6. Molenaar, R.J., Maciejewski, J.P., Wilmink, J.W., and van Noorden, C.J.F. 2018. Wild-type and mutated IDH1/2 enzymes and therapy responses. Oncogene 37:1949–1960.

7. Verhaak, R.G., Hoadley, K.A., Purdom, E., Wang, V., Qi, Y., Wilkerson, M.D., Miller, C.R., Ding, L., Golub, T., Mesirov, J.P. et al 2010. Integrated genomic analysis identifies clinically relevant subtypes of glioblastoma characterized by abnormalities in PDGFRA, IDH1, EGFR, and NF1. Cancer Cell 17:98–110.

8. Ichimura, K., Narita, Y., and Hawkins, C.E. 2015. Diffusely infiltrating astrocytomas: pathology, molecular mechanisms and markers. Acta Neuropathol. 129:789–808.

9. Avila, J., de Barreda, E.G., Fuster-Matanzo, A., Simon, D., Llorens-Martin, M., Engel, T., Lucas, J.J., Diaz-Hernandez, M., and Hernandez, F. 2012. Looking for novel functions of tau. Biochem. Soc. Trans. 40:653–655.

10. Barreda, E.G., and Avila, J. 2011. Tau regulates the subcellular localization of calmodulin. Biochem. Biophys. Res. Commun. 408:500–504.

11. Li, H., Duan, Z.W., Xie, P., Liu, Y.R., Wang, W.C., Dou, S.X., and Wang, P.Y. 2012. Effects of paclitaxel on EGFR endocytic trafficking revealed using quantum dot tracking in single cells. PLoS. One. 7:e45465.

12. Lehrer, S. 2010. Glioblastoma and dementia may share a common cause. Med. Hypotheses 75:67–68.

13. Nunez, F.J., Mendez, F.M., Kadiyala, P., Alghamri, M.S., Savelieff, M.G., Garcia-Fabiani, M.B., Haase, S., Koschmann, C., Calinescu, A.A., Kamran, N. et al 2019. IDH1-R132H acts as a tumor suppressor in glioma via epigenetic up-regulation of the DNA damage response. Sci. Transl. Med. 11.

14. Flavahan, W.A., Drier, Y., Liau, B.B., Gillespie, S.M., Venteicher, A.S., Stemmer-Rachamimov, A.O., Suva, M.L., and Bernstein, B.E. 2016. Insulator dysfunction and oncogene activation in IDH mutant gliomas. Nature 529:110–114.

15. Caillet-Boudin, M.L., Buee, L., Sergeant, N., and Lefebvre, B. 2015. Regulation of human MAPT gene expression. Mol. Neurodegener. 10:28.

16. Perez, M., Santa-Maria, I., Gomez de, B.E., Zhu, X., Cuadros, R., Cabrero, J.R., Sanchez-Madrid, F., Dawson, H.N., Vitek, M.P., Perry, G. et al 2009. Tau--an inhibitor of deacetylase HDAC6 function. J. Neurochem. 109:1756–1766.

17. Gao, Y.S., Hubbert, C.C., and Yao, T.P. 2010. The microtubule-associated histone deacetylase 6 (HDAC6) regulates epidermal growth factor receptor (EGFR) endocytic trafficking and degradation. J. Biol. Chem. 285:11219–11226.

18. Zhang, B., Carroll, J., Trojanowski, J.Q., Yao, Y., Iba, M., Potuzak, J.S., Hogan, A.M., Xie, S.X., Ballatore, C., Smith, A.B., III et al 2012. The microtubule-stabilizing agent, epothilone D, reduces axonal dysfunction, neurotoxicity, cognitive deficits, and Alzheimer-like pathology in an interventional study with aged tau transgenic mice. J. Neurosci. 32:3601–3611.

19. Bhat, K.P., Balasubramaniyan, V., Vaillant, B., Ezhilarasan, R., Hummelink, K., Hollingsworth, F., Wani, K., Heathcock, L., James, J.D., Goodman, L.D. et al 2013. Mesenchymal differentiation mediated by NF-kappaB promotes radiation resistance in glioblastoma. Cancer Cell 24:331–346.

20. Pozo, N., Zahonero, C., Fernandez, P., Linares, J.M., Ayuso, A., Hagiwara, M., Perez, A., Ricoy, J.R., Hernandez-Lain, A., Sepulveda, J.M. et al 2013. Inhibition of DYRK1A destabilizes EGFR and reduces EGFR-dependent glioblastoma growth. J Clin Invest 123:2475–2487.

21. Lu, F., Chen, Y., Zhao, C., Wang, H., He, D., Xu, L., Wang, J., He, X., Deng, Y., Lu, E.E. et al 2016. Olig2-Dependent Reciprocal Shift in PDGF and EGF Receptor Signaling Regulates Tumor Phenotype and Mitotic Growth in Malignant Glioma. Cancer Cell 29:669–683.

22. Garcia-Romero, N., Gonzalez-Tejedo, C., Carrion-Navarro, J., Esteban-Rubio, S., Rackov, G., Rodriguez-Fanjul, V., Oliver-De La Cruz, J., Prat-Acin, R., Peris-Celda, M., Blesa, D. et al 2016. Cancer stem cells from human glioblastoma resemble but do not mimic original tumors after in vitro passaging in serum-free media. Oncotarget. 7:65888–65901.

23. Bougnaud, S., Golebiewska, A., Oudin, A., Keunen, O., Harter, P.N., Mader, L., Azuaje, F., Fritah, S., Stieber, D., Kaoma, T. et al 2016. Molecular crosstalk between tumour and brain parenchyma instructs histopathological features in glioblastoma. Oncotarget. 7:31955–31971.

24. Bergers, G., and Song, S. 2005. The role of pericytes in blood-vessel formation and maintenance. Neuro. Oncol. 7:452–464.

25. Scully, S., Francescone, R., Faibish, M., Bentley, B., Taylor, S.L., Oh, D., Schapiro, R., Moral, L., Yan, W., and Shao, R. 2012. Transdifferentiation of glioblastoma stem-like cells into mural cells drives vasculogenic mimicry in glioblastomas. J. Neurosci. 32:12950–12960.

26. Cheng, L., Huang, Z., Zhou, W., Wu, Q., Donnola, S., Liu, J.K., Fang, X., Sloan, A.E., Mao, Y., Lathia, J.D. et al 2013. Glioblastoma stem cells generate vascular pericytes to support vessel function and tumor growth. Cell 153:139–152.

27. Zhou, W., Chen, C., Shi, Y., Wu, Q., Gimple, R.C., Fang, X., Huang, Z., Zhai, K., Ke, S.Q., Ping, Y.F. et al 2017. Targeting Glioma Stem Cell-Derived Pericytes Disrupts the Blood-Tumor Barrier and Improves Chemotherapeutic Efficacy. Cell Stem Cell 21:591–603.

28. Hardee, M.E., and Zagzag, D. 2012. Mechanisms of glioma-associated neovascularization. Am. J. Pathol. 181:1126–1141.

29. Park, J.S., Kim, I.K., Han, S., Park, I., Kim, C., Bae, J., Oh, S.J., Lee, S., Kim, J.H., Woo, D.C. et al 2016. Normalization of Tumor Vessels by Tie2 Activation and Ang2 Inhibition Enhances Drug Delivery and Produces a Favorable Tumor Microenvironment. Cancer Cell 30:953–967.

30. Simonavicius, N., Robertson, D., Bax, D.A., Jones, C., Huijbers, I.J., and Isacke, C.M. 2008. Endosialin (CD248) is a marker of tumor-associated pericytes in high-grade glioma. Mod. Pathol. 21:308–315.

31. Buckner, J.C., Chakravarti, A., and Curran, W.J., Jr. 2016. Radiation plus Chemotherapy in Low-Grade Glioma. N. Engl. J. Med. 375:490–491.

32. Cairncross, J.G., Wang, M., Jenkins, R.B., Shaw, E.G., Giannini, C., Brachman, D.G., Buckner, J.C., Fink, K.L., Souhami, L., Laperriere, N.J. et al 2014. Benefit from procarbazine, lomustine, and vincristine in oligodendroglial tumors is associated with mutation of IDH. J. Clin. Oncol. 32:783–790.

33. Amankulor, N.M., Kim, Y., Arora, S., Kargl, J., Szulzewsky, F., Hanke, M., Margineantu, D.H., Rao, A., Bolouri, H., Delrow, J. et al 2017. Mutant IDH1 regulates the tumor-associated immune system in gliomas. Genes Dev. 31:774–786.

34. Yao, Y., Chan, A.K., Qin, Z.Y., Chen, L.C., Zhang, X., Pang, J.C., Li, H.M., Wang, Y., Mao, Y., Ng, H.K. et al 2013. Mutation analysis of IDH1 in paired gliomas revealed IDH1 mutation was not associated with malignant progression but predicted longer survival. PLoS. One. 8:e67421.

35. Johnson, B.E., Mazor, T., Hong, C., Barnes, M., Aihara, K., McLean, C.Y., Fouse, S.D., Yamamoto, S., Ueda, H., Tatsuno, K. et al 2014. Mutational analysis reveals the origin and therapy-driven evolution of recurrent glioma. Science 343:189–193.

36. Watanabe, T., Nobusawa, S., Kleihues, P., and Ohgaki, H. 2009. IDH1 mutations are early events in the development of astrocytomas and oligodendrogliomas. Am. J. Pathol. 174:1149–1153.

37. Bardella, C., Al-Dalahmah, O., Krell, D., Brazauskas, P., Al-Qahtani, K., Tomkova, M., Adam, J., Serres, S., Lockstone, H., Freeman-Mills, L. et al 2016. Expression of Idh1(R132H) in the Murine Subventricular Zone Stem Cell Niche Recapitulates Features of Early Gliomagenesis. Cancer Cell 30:578–594.

38. Philip, B., Yu, D.X., Silvis, M.R., Shin, C.H., Robinson, J.P., Robinson, G.L., Welker, A.E., Angel, S.N., Tripp, S.R., Sonnen, J.A. et al 2018. Mutant IDH1 Promotes Glioma Formation In Vivo. Cell Rep. 23:1553–1564.

39. Rohle, D., Popovici-Muller, J., Palaskas, N., Turcan, S., Grommes, C., Campos, C., Tsoi, J., Clark, O., Oldrini, B., Komisopoulou, E. et al 2013. An inhibitor of mutant IDH1 delays growth and promotes differentiation of glioma cells. Science 340:626–630.

40. Golub, D., Iyengar, N., Dogra, S., Wong, T., Bready, D., Tang, K., Modrek, A.S., and Placantonakis, D.G. 2019. Mutant Isocitrate Dehydrogenase Inhibitors as Targeted Cancer Therapeutics. Front Oncol. 9:417.

41. Molenaar, R.J., Botman, D., Smits, M.A., Hira, V.V., van Lith, S.A., Stap, J., Henneman, P., Khurshed, M., Lenting, K., Mul, A.N. et al 2015. Radioprotection of IDH1-Mutated Cancer Cells by the IDH1-Mutant Inhibitor AGI-5198. Cancer Res. 75:4790–4802.

42. Khurshed, M., Aarnoudse, N., Hulsbos, R., Hira, V.V.V., van Laarhoven, H.W.M., Wilmink, J.W., Molenaar, R.J., and van Noorden, C.J.F. 2018. IDH1-mutant cancer cells are sensitive to cisplatin and an IDH1-mutant inhibitor counteracts this sensitivity. FASEB J. fj201800547R.

43. Wu, X., Sooman, L., Lennartsson, J., Bergstrom, S., Bergqvist, M., Gullbo, J., and Ekman, S. 2013. Microtubule inhibition causes epidermal growth factor receptor inactivation in oesophageal cancer cells. Int. J. Oncol. 42:297–304.

44. Zahonero, C., and Sanchez-Gomez, P. 2014. EGFR-dependent mechanisms in glioblastoma: towards a better therapeutic strategy. Cell Mol. Life Sci.

45. Grandal, M.V., Zandi, R., Pedersen, M.W., Willumsen, B.M., van, D.B., and Poulsen, H.S. 2007. EGFRvIII escapes down-regulation due to impaired internalization and sorting to lysosomes. Carcinogenesis 28:1408–1417.

46. Bonavia, R., Inda, M.M., Vandenberg, S., Cheng, S.Y., Nagane, M., Hadwiger, P., Tan, P., Sah, D.W., Cavenee, W.K., and Furnari, F.B. 2012. EGFRvIII promotes glioma angiogenesis and growth through the NF-kappaB, interleukin-8 pathway. Oncogene 31:4054–4066.

47. Gargini, R., Escoll, M., Garcia, E., Garcia-Escudero, R., Wandosell, F., and Anton, I.M. 2016. WIP Drives Tumor Progression through YAP/TAZ-Dependent Autonomous Cell Growth. Cell Rep. 17:1962–1977.

48. Ferraiuolo, M., Pulito, C., Finch-Edmondson, M., Korita, E., Maidecchi, A., Donzelli, S., Muti, P., Serra, M., Sudol, M., Strano, S. et al 2018. Agave negatively regulates YAP and TAZ transcriptionally and post-translationally in osteosarcoma cell lines. Cancer Lett. 433:18–32.

49. Lai, A., Kharbanda, S., Pope, W.B., Tran, A., Solis, O.E., Peale, F., Forrest, W.F., Pujara, K., Carrillo, J.A., Pandita, A. et al 2011. Evidence for sequenced molecular evolution of IDH1 mutant glioblastoma from a distinct cell of origin. J. Clin. Oncol. 29:4482–4490.

50. Unruh, D., Schwarze, S.R., Khoury, L., Thomas, C., Wu, M., Chen, L., Chen, R., Liu, Y., Schwartz, M.A., Amidei, C. et al 2016. Mutant IDH1 and thrombosis in gliomas. Acta Neuropathol. 132:917–930.

51. Kickingereder, P., Sahm, F., Radbruch, A., Wick, W., Heiland, S., Deimling, A., Bendszus, M., and Wiestler, B. 2015. IDH mutation status is associated with a distinct hypoxia/angiogenesis transcriptome signature which is non-invasively predictable with rCBV imaging in human glioma. Sci. Rep. 5:16238.

52. Missiaen, R., Mazzone, M., and Bergers, G. 2018. The reciprocal function and regulation of tumor vessels and immune cells offers new therapeutic opportunities in cancer. Semin. Cancer Biol. 52:107–116.

53. Pallas-Bazarra, N., Jurado-Arjona, J., Navarrete, M., Esteban, J.A., Hernandez, F., Avila, J., and Llorens-Martin, M. 2016. Novel function of Tau in regulating the effects of external stimuli on adult hippocampal neurogenesis. EMBO J. 35:1417–1436.

54. Podtelezhnikov, A.A., Tanis, K.Q., Nebozhyn, M., Ray, W.J., Stone, D.J., and Loboda, A.P. 2011. Molecular insights into the pathogenesis of Alzheimer’s disease and its relationship to normal aging. PLoS. One. 6:e29610.

55. Kisler, K., Nelson, A.R., Montagne, A., and Zlokovic, B.V. 2017. Cerebral blood flow regulation and neurovascular dysfunction in Alzheimer disease. Nat. Rev. Neurosci. 18:419–434.

56. Sagare, A.P., Bell, R.D., Zhao, Z., Ma, Q., Winkler, E.A., Ramanathan, A., and Zlokovic, B.V. 2013. Pericyte loss influences Alzheimer-like neurodegeneration in mice. Nat. Commun. 4:2932.

57. Semiond, D., Sidhu, S.S., Bissery, M.C., and Vrignaud, P. 2013. Can taxanes provide benefit in patients with CNS tumors and in pediatric patients with tumors? An update on the preclinical development of cabazitaxel. Cancer Chemother. Pharmacol. 72:515–528.

58. Smoter, M., Bodnar, L., Duchnowska, R., Stec, R., Grala, B., and Szczylik, C. 2011. The role of Tau protein in resistance to paclitaxel. Cancer Chemother. Pharmacol. 68:553–557.

59. Ferron, S.R., Andreu-Agullo, C., Mira, H., Sanchez, P., Marques-Torrejon, M.A., and Farinas, I. 2007. A combined ex/in vivo assay to detect effects of exogenously added factors in neural stem cells. Nat. Protoc. 2:849–859.

