## Supplementary data for "IDH-Tau-EGFR triad defines the neovascular landscape of diffuse gliomas by controlling mesenchymal differentiation"

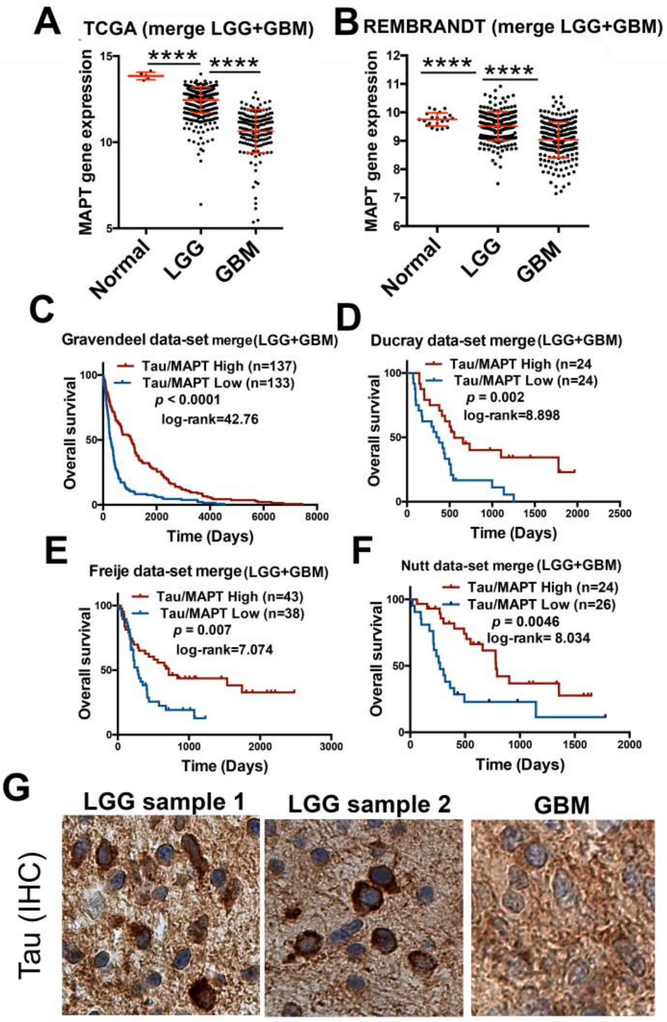


**Fig. S1. Association of Tau levels with the clinical pathology of diffuse gliomas.** (**A** and **B**) Analysis of *Tau (MAPT)* mRNA expression by RNAseq in diffuse glioma patients compared to normal tissue, using the TCGA (n=692) (A) and the Rembrandt (n=432) (B) cohorts. Tumors were grouped according to the WHO classification (histological type). (**C-F**) Kaplan-Meier overall survival curves of patients from the Gravendeel (n=276) (**C**); the Ducray (n=48) (**D**); the Freije (n=81) (**E**) and the Nutt (n=60) (**F**) cohorts. Patients in each cohort were stratified into 2 groups based on high and low *Tau (MAPT)* expression values. (**G**) Representative pictures of the IHC Tau staining of several gliomas. ****, $p \leq 0.0001$.

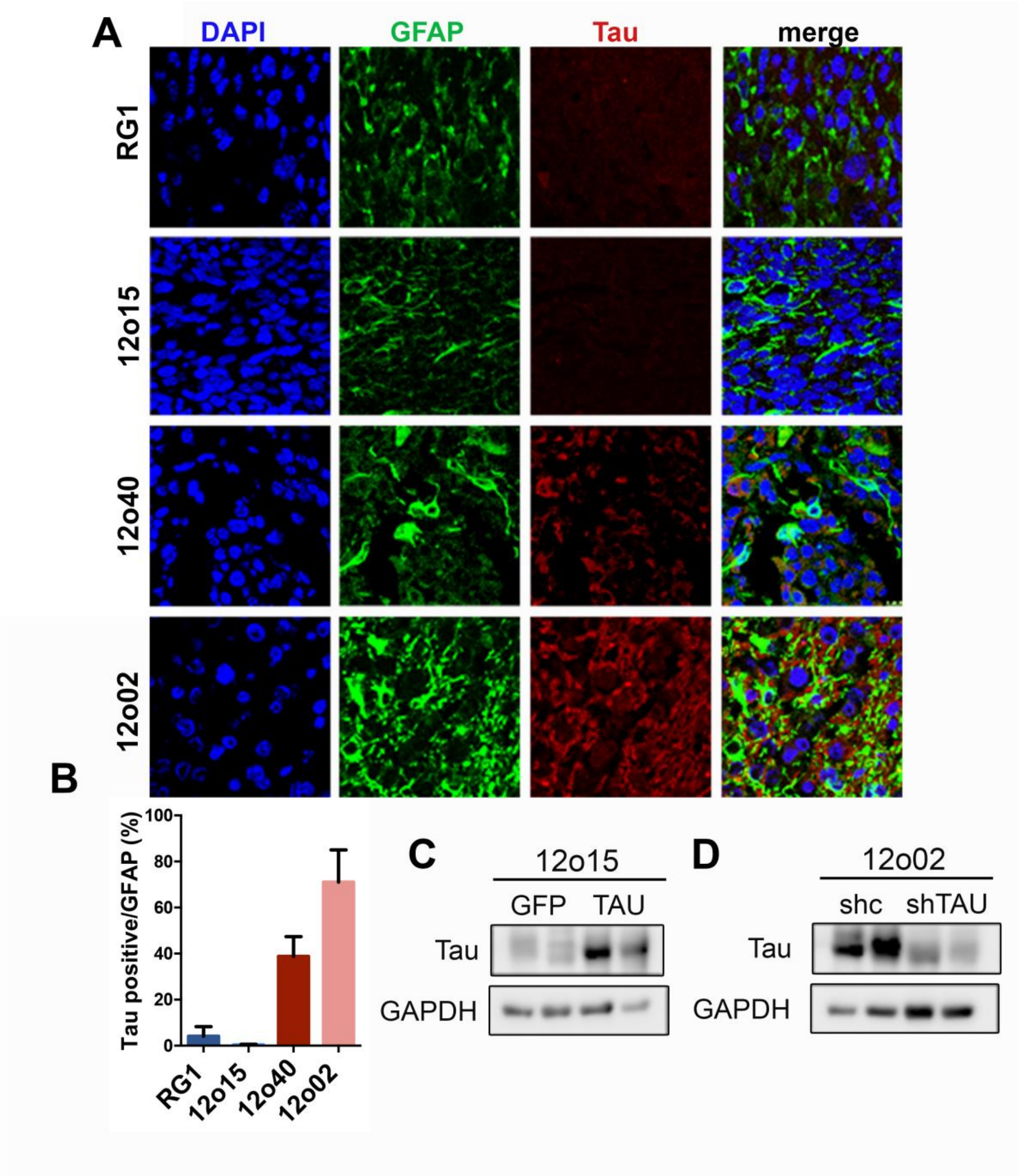


**Fig. S2.** (**A**) Representative images of Tau and GFAP by IF co-staining of tumors from Fig. 2H. (**B**) Quantification of the co-staining of Tau and GFAP in A. (**C-D**) WB analysis and quantification of Tau in the tumors from Fig. 2J (C) and Fig. 2K (D). GAPDH levels were used for normalization.

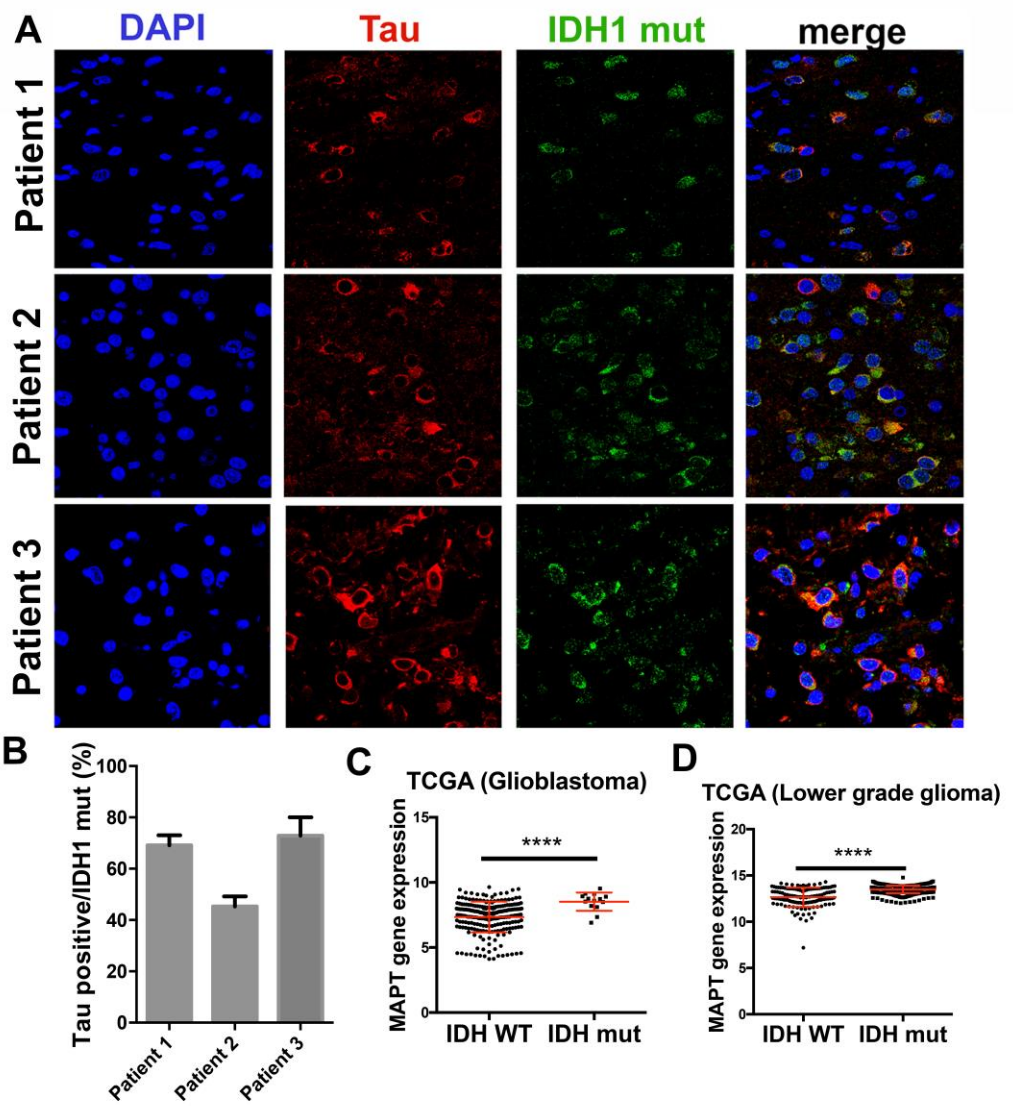


**Fig. S3. Tau expression correlates with the presence of IDH mutations.** (**A**) Representative images of Tau and IDH1mut IF co-staining of tumors from three different patients. (**B**) Quantification of the co-staining of Tau and IDH1mut in A. (**C** and **D**) Analysis of *Tau (MAPT)* mRNA expression by RNAseq in the GBM (C) and the LGG (D) TCGA cohorts, grouped based on the presence of IDH mutations (n=692). ****, $p \leq 0.0001$.

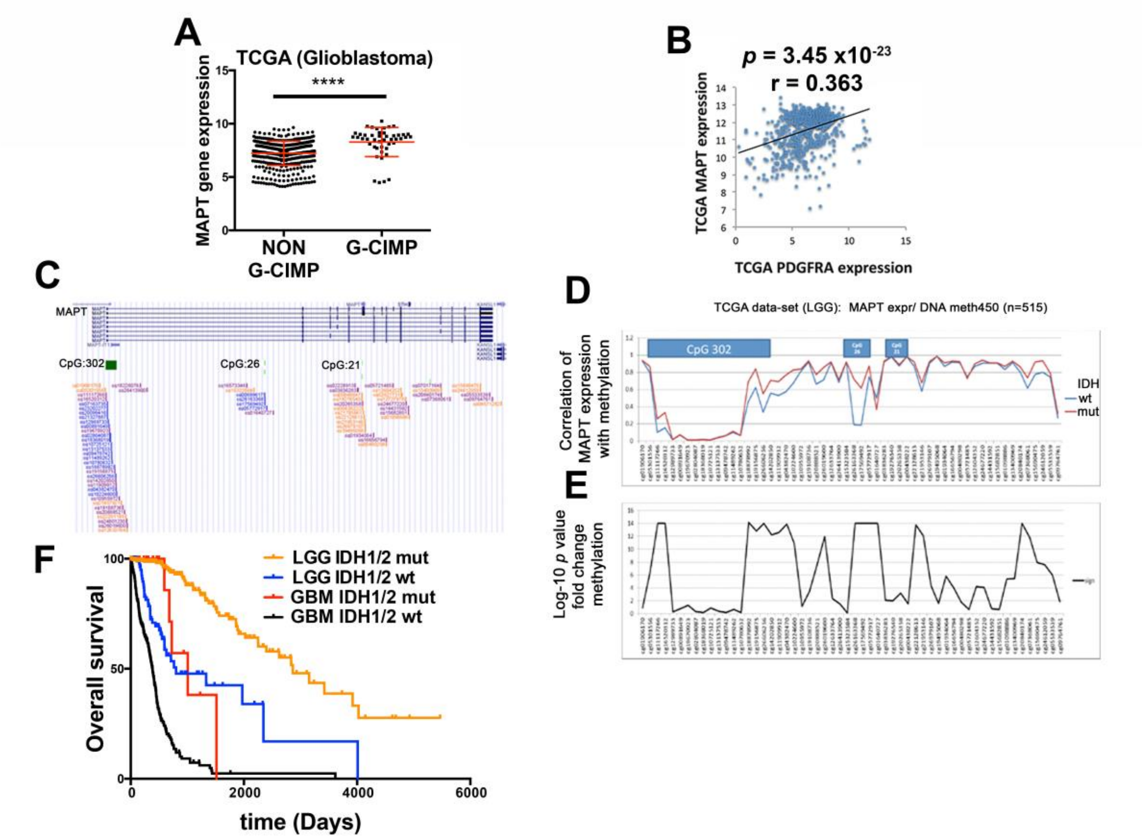


**Fig. S4. The expression of Tau is associated with the IDH mutant methylation phenotype.** (**A**) Analysis of *Tau (MAPT)* mRNA expression in GBM according to the level of CpG methylation: low-level methylator phenotype (NON G-CIMP) or high-level methylator phenotype (G-CIMP). (**B**) Correlation of the expression of *Tau (MAPT)* with that of *PDGFRA* (n=702) using the TCGA-merge (LGG+GBM) dataset. (**C**) Organization of the possible CpG islets on the promoter zone of the *Tau* (MAPT) gene using the methylation probes by genome browser. (**D** and **E**) Analysis of the fold change CpG methylation in 515 patients with LGG. The blue line shows the levels of methylation in gliomas with wildtype *IDH* and in red the level of methylation in gliomas with mutations in *IDH*. (**F**) Kaplan-Meier overall survival curves of LGG (n=451) and GBM (n=299) patients from the TCGA cohort, stratified based on the status of IDH1/2. ****, $p \leq 0.0001$.

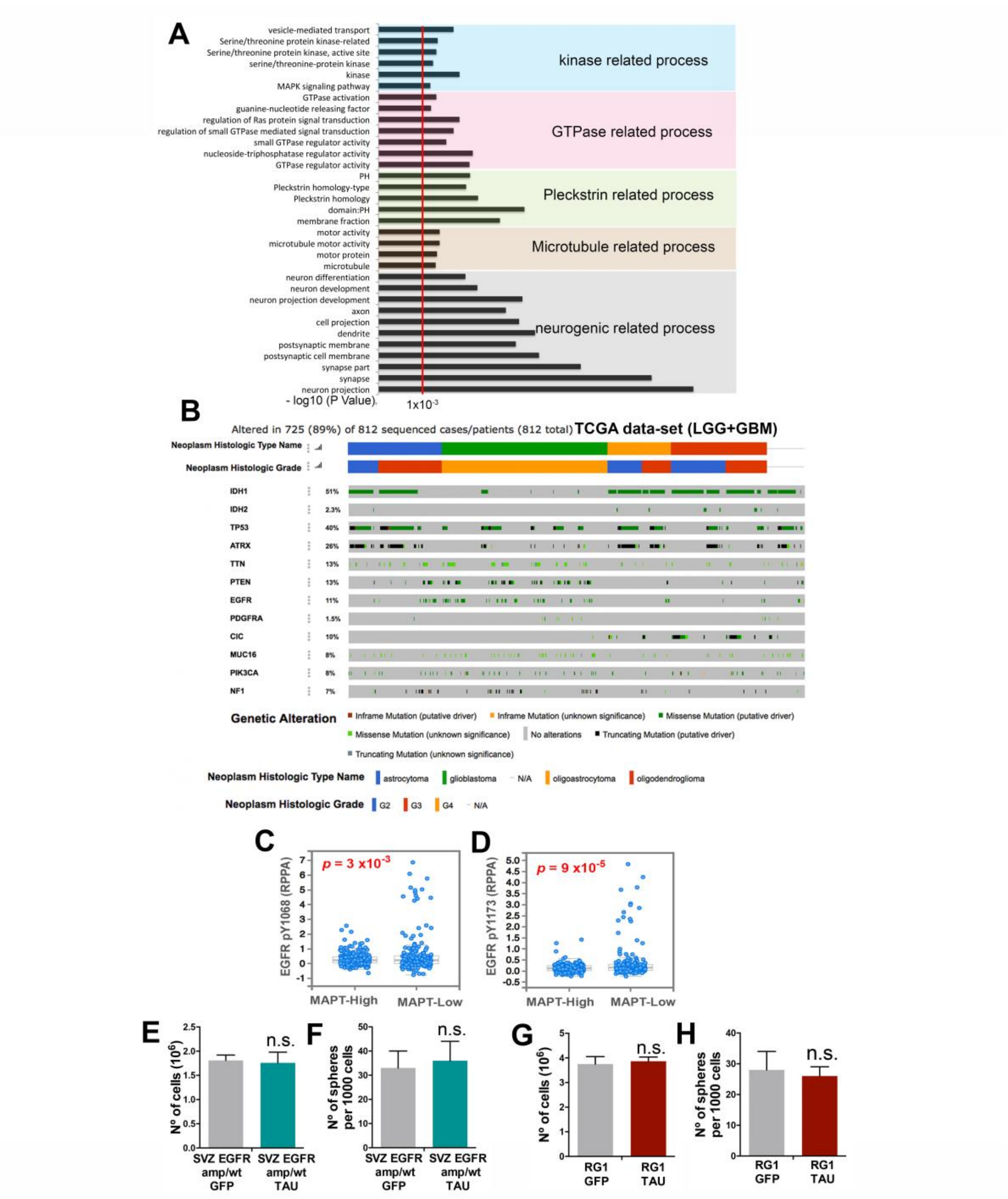


**Fig. S5. Association of Tau function with the EGFR pathway in gliomas.** (**A**) Top enriched Gene Ontology (GO) biological process for the cluster of 500 genes that are positively co-expressed with Tau (MAPT) in gliomas. We used the LGG+GBM merge cohort and the DAVID gene ontology program. (**B**) Histogram showing the non-silent somatic mutations in genes commonly modified in diffuse gliomas grouped according to the WHO classification (histological type and grade). (**C**) Analysis of levels of Phospho-Tyr1068-EGFR and Phospho-Tyr1173-EGFR in a cohort of 244 patients with GBM (TCGA-GBM dataset) according to high or low levels of Tau (MAPT). (**E** to **H**) Quantification of the number of cells (**E** and **G**) and spheres (**F** and **H**) in SVZ-EGFRamp/wt (**E** and **F**) and RG (**G** and **H**) cells after the overexpression of GFP or Tau (n = 3). n.s., non-significant.

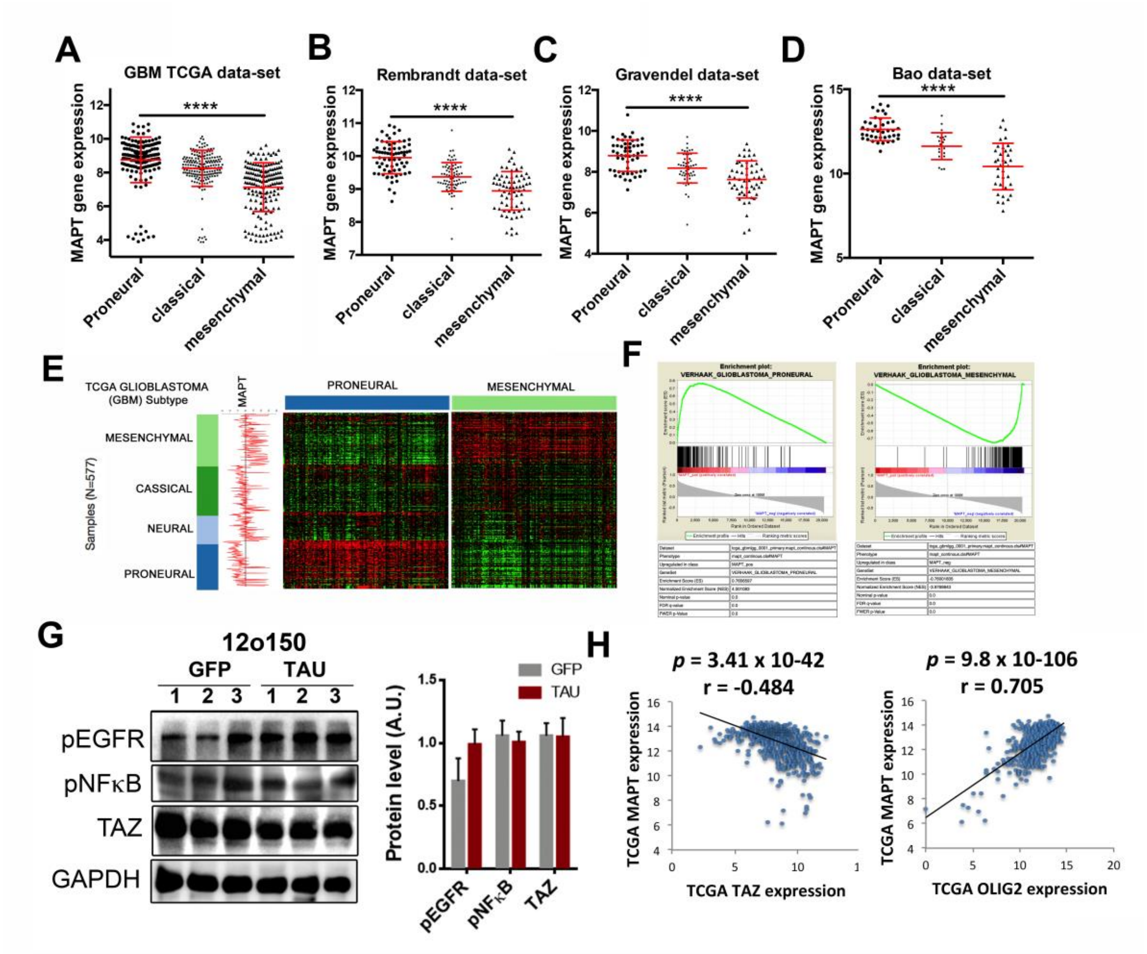


**Fig. S6. Association of Tau expression with the GBM subtypes and the NF-kB-TAZ axis.** (**A** to **D**) Analysis of *Tau (MAPT)* mRNA expression in the different GBM subtypes using the TCGA (n=528) (A), the Rembrandt (n=219) (B), the Gravendeel (n=159) (C) and the Bao (n=100) (D) data sets. (**E**) Heatmap of PN and MES gene expression signature depending on the levels of expression of Tau and analysis of mRNA Tau levels according to GBM subtypes. (**F**) GSEA enrichment plot analysis using Tau gene expression values as template and PN or MES signatures. (**G**) WB analysis of phospho-EGFR, phosphor-NF-κB and TAZ in 12o150 xenografts expressing either GFP or Tau. GADPH levels were used for normalization. Quantification is shown on the right. (**H**) Scatter plots showing the correlation between the expression of Tau (MAPT) and TAZ genes (left) or Tau (MAPT) and OLIG2 genes (right) across 703 glioma samples. ****, $p \leq 0.0001$.

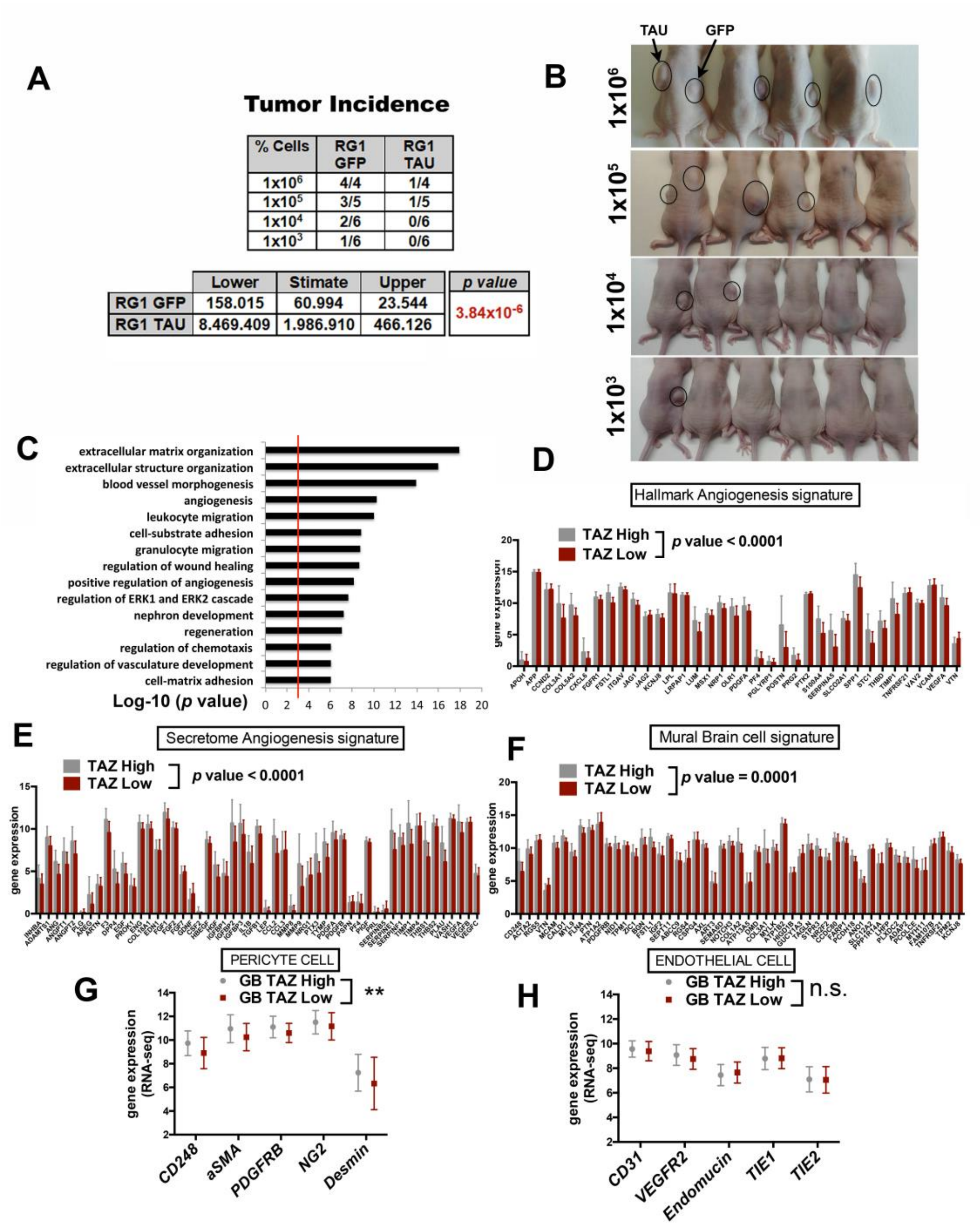


**Fig. S7. Vascular phenotypes associated with TAZ in gliomas.** (**A** and **B**) Subcutaneous tumor growth assay (limited dilution) of RG1 cells after overexpression of GFP or Tau. The statistical analysis is shown on the bottom and the image of the animals before tumor dissection is shown in B. (**C**) The top 15 gene ontology (GO) terms associated with cluster from the 500 genes that correlate positively with the expression of TAZ in TCGA-LGG+GBM dataset. GO terms were ranked by p value. (**D** to **F**) Angiogenesis (**D**), angiogenic secretome (**E**) and mural brain cell (**F**) signatures in gliomas of the TCGA-LGG+GBM-merge data-set. Tumors were classified as High or Low Taz-expressing gliomas. (**G** and **H**) Analysis of the expression (RNAseq) of the 5 most significant genes associated with pericytes (G) or endothelial cells (H); signature including the most relevant genes for biological processes associated with vasculogenesis, angiogenesis and secretion in (C-F). ** $p \leq 0.01$. n.s. non significant.

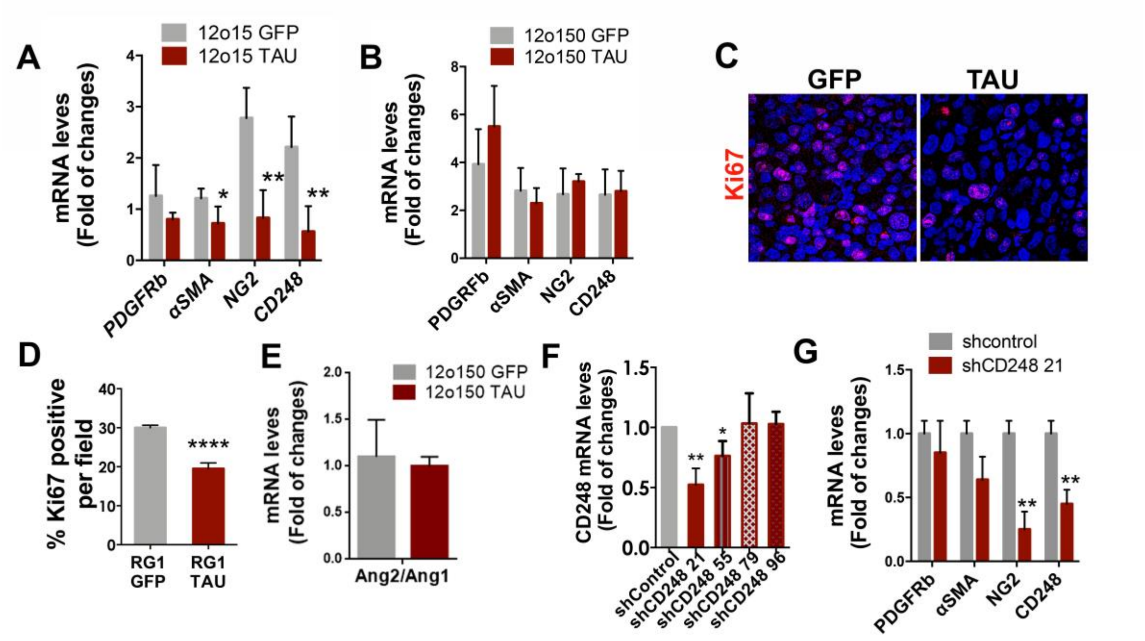


**Fig. S8. Implication of the tumor-derived-pericytes on the glioma vasculature and growth.** (**A** and **B**) qRT-PCR analysis of pericytic–related genes (using human-specific primers) in 12o15 (A) and 12o150 (B) xenografts expressing GFP or Tau. Mouse cDNA was used as a negative control. The expression of *HPRT* was used for normalization**.** (**C** and **D**) Representative images of Ki67 IF staining of sections from RG1 xenografts expressing GFP or Tau (C) and quantification of the number of Ki67 positive cells per field (D). (**E**) Ratio of *Ang2/Ang1* expression measured by qRT-PCR in 12o150 tumors. The expression of HPRT was used for normalization. (**F**) qRT-PCR analysis of *CD248* levels after transduction of different shRNAs against the *CD248* gene in RG1 cells. The expression of *HPRT* was used for normalization. (**G**) qRT-PCR quantification of 4 pericyte markers: *CD248, NG2, αSMA* and *PDGFRβ*, after the transduction of shcontrol or shCD248 21 in the RG1 line. The expression of *HPRT* was used for normalization. *, *p* ≤0.05; **, *p* ≤0.01; ****, *p* ≤0.0001.

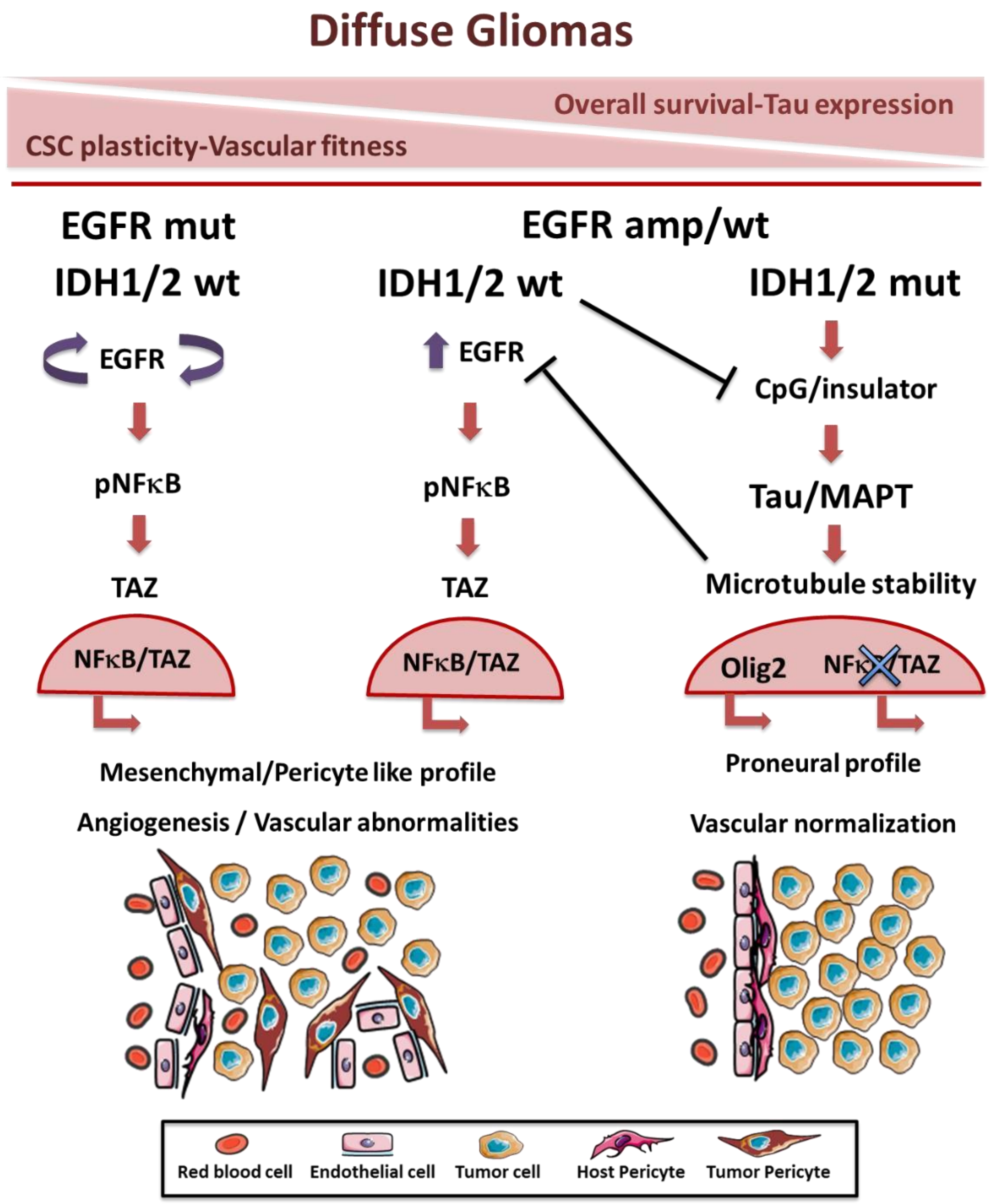


**Fig. S9. Integrated stratification of diffuse gliomas based on the status of *EGFR* and *IDH* and the expression of Tau.** EGFRmut and EGFRamp glioma cells generate pericytes (glioma-derived-pericytes, GDPs) with distinct vascular capacities. The GDPs are the master regulators of the vascular landscape, defining the leakiness of the BBB and the proliferative capacity of the tumors. In IDHmutant tumors, Tau expression is epigenetically induced and impairs the appearance of GDPs by inhibiting the EGFRamp-NF B-TAZ axis status.

**Supplementary Table 1. Human samples**

| Sample | Hospital | Diagnosis | Grade | Tau IHC | IDH | ATRX |
|---|---|---|---|---|---|---|
| 1 | H12O | Glioblastoma | IV | 0 | wt | wt |
| 2 | H12O | Glioblastoma | IV | 0 | wt | wt |
| 3 | H12O | Glioblastoma | IV | 1 | wt | wt |
| 4 | H12O | Glioblastoma | IV | 0 | wt | wt |
| 5 | H12O | Glioblastoma | IV | 0 | wt | wt |
| 6 | H12O | Glioblastoma | IV | 2 | wt | wt |
| 7 | H12O | Glioblastoma | IV | 3 | wt | wt |
| 8 | H12O | Glioblastoma | IV | 3 | wt | wt |
| 9 | H12O | Glioblastoma | IV | 1 | wt | wt |
| 10 | H12O | Glioblastoma | IV | 1 | wt | wt |
| 11 | H12O | Glioblastoma | IV | 0 | wt | wt |
| 12 | H12O | Anaplastic Astrocytoma | nd | 3 | mut | wt |
| 13 | H12O | Glioblastoma | IV | 1 | wt | wt |
| 14 | H12O | Glioblastoma | IV | 0 | wt | mut |
| 15 | H12O | Glioblastoma | IV | 1 | wt | mut |
| 16 | H12O | Glioblastoma | IV | 2 | wt | wt |
| 17 | H12O | Glioblastoma | IV | 3 | wt | wt |
| 18 | H12O | Glioblastoma | IV | 2 | wt | mut |
| 19 | H12O | Glioblastoma | IV | 0 | wt | wt |
| 20 | H12O | Glioblastoma | IV | 0 | wt | wt |
| 21 | H12O | Glioblastoma | IV | 1 | wt | wt |
| 22 | H12O | Glioblastoma | IV | 0 | wt | wt |
| 23 | H12O | Glioblastoma | IV | 1 | wt | wt |
| 24 | H12O | Glioblastoma | IV | 0 | wt | wt |
| 25 | H12O | Glioblastoma | IV | 1 | wt | mut |
| 26 | H12O | Glioblastoma | IV | 3 | wt | wt |
| 27 | H12O | Glioblastoma | IV | 1 | wt | wt |
| 28 | H12O | Glioblastoma | IV | 0 | wt | wt |
| 29 | H12O | Glioblastoma | IV | 0 | wt | wt |
| 30 | H12O | Glioblastoma | IV | 1 | wt | wt |
| 31 | H12O | Glioblastoma | IV | 1 | wt | wt |
| 32 | H12O | Glioblastoma | IV | 0 | wt | wt |
| 33 | H12O | Glioblastoma | IV | 0 | wt | wt |
| 34 | H12O | Oligodendroglioma | nd | 1 | mut | wt |
| 35 | H12O | Oligodendroglioma | III | 2 | mut | wt |
| 36 | H12O | Diffuse mindline glioma | nd | 2 | wt | mut |
| 37 | H12O | Anaplastic Astrocytoma | III | 0 | wt | wt |
| 38 | H12O | Astrocytoma | II | 1 | wt | nd |
| 39 | H12O | Anaplastic Astrocytoma | III | 3 | mut | mut |
| 40 | H12O | Anaplastic Astrocytoma | III | 3 | mut | mut |
| 41 | H12O | Diffuse Astrocytoma | II | 3 | mut | mut |

| | | | | | | |
|---|---|---|---|---|---|---|
| 42 | H12O | Anaplastic Astrocytoma | III | 3 | wt | mut |
| 43 | H12O | Diffuse Astrocytoma | II | 3 | wt | wt |
| 44 | H12O | Anaplastic Astrocytoma | III | 1 | mut | mut |
| 45 | H12O | Anaplastic Astrocytoma | III | 2 | mut | nd |
| 46 | H12O | Anaplastic Astrocytoma | III | 3 | wt | wt |
| 47 | H12O | Anaplastic Astrocytoma | III | 2 | mut | mut |
| 48 | H12O | Anaplastic Astrocytoma | III | 3 | wt | wt |
| 49 | H12O | Anaplastic Astrocytoma | III | 0 | mut | nd |
| 50 | H12O | Astrocytoma | II | 2 | mut | nd |
| 51 | H12O | Anaplastic Astrocytoma | III | 3 | wt | mut |
| 52 | H12O | Glioblastoma | IV | 3 | mut | mut |
| 53 | H12O | Astrocytoma | II | 2 | mut | mut |
| 54 | H12O | Anaplastic Astrocytoma | III | 3 | mut | mut |
| 55 | H12O | Anaplastic Astrocytoma | III | 1 | wt | wt |
| 56 | H12O | Anaplastic Astrocytoma | III | 0 | mut | mut |
| 57 | H12O | Anaplastic Astrocytoma | III | 3 | mut | mut |
| 58 | H12O | Anaplastic Astrocytoma | III | 3 | mut | nd |
| 59 | H12O | Anaplastic Astrocytoma | III | 3 | mut | nd |
| 60 | H12O | Anaplastic Astrocytoma | III | 1 | mut | mut |
| 61 | H12O | Anaplastic Astrocytoma | III | 2 | mut | mut |
| 62 | H12O | Astrocytoma | II | 3 | mut | mut |
| 63 | H12O | Diffuse Astrocytoma | II | 2 | mut | mut |
| 64 | H12O | Anaplastic Astrocytoma | III | 3 | mut | mut |
| 65 | H12O | Astrocytoma | II | 2 | wt | nd |
| 66 | H12O | Diffuse Astrocytoma | II | 3 | mut | mut |
| 67 | H12O | Anaplastic Astrocytoma | III | 1 | mut | mut |
| 68 | H12O | Anaplastic Astrocytoma | III | 1 | wt | nd |
| 69 | H12O | Anaplastic Astrocytoma | III | 3 | wt | nd |
| 70 | La Fe | Glioblastoma | IV | 0 | nd | nd |
| 71 | La Fe | Glioblastoma | IV | 0 | nd | nd |
| 72 | La Fe | Glioblastoma | IV | 0 | nd | nd |
| 73 | La Fe | Glioblastoma | IV | 0 | nd | nd |
| 74 | La Fe | Glioblastoma | IV | 0 | nd | nd |
| 75 | La Fe | Glioblastoma | IV | 0 | nd | nd |
| 76 | La Fe | Glioblastoma | IV | 0 | nd | nd |
| 77 | La Fe | Glioblastoma | IV | 0 | nd | nd |
| 78 | La Fe | Glioblastoma | IV | 0 | nd | nd |
| 79 | La Fe | Glioblastoma | IV | 0 | nd | nd |

| | | | | | | |
|---|---|---|---|---|---|---|
| 80 | La Fe | Glioblastoma | IV | 0 | nd | nd |
| 81 | La Fe | Oligodendroglioma | nd | 1 | nd | nd |
| 82 | La Fe | Glioblastoma | IV | 2 | nd | nd |
| 83 | HGM | Glioblastoma | IV | 1 | nd | nd |
| 84 | HGM | Oligoastrocytoma | II | 2 | mut | nd |
| 85 | HGM | Oligodendroglioma | II | 3 | mut | nd |
| 86 | La Fe | Glioblastoma | IV | 0 | nd | nd |
| 87 | La Fe | Glioblastoma | IV | 2 | nd | nd |
| 88 | La Fe | Glioblastoma | IV | 1 | nd | nd |
| 89 | La Fe | Glioblastoma | IV | 0 | nd | nd |
| 90 | La Fe | Glioblastoma | IV | 0 | nd | nd |
| 91 | La Fe | Glioblastoma | IV | 0 | nd | nd |
| 92 | La Fe | Anaplastic Oligoastrocytoma | III | 2 | nd | nd |
| 93 | La Fe | Anaplastic Oligoastrocytoma | III | 3 | mut | nd |
| 94 | La Fe | Diffuse Glioma | II | 3 | mut | nd |
| 95 | La Fe | Glioblastoma | IV | 2 | wt | nd |
| 96 | La Fe | Glioblastoma | IV | 0 | wt | nd |
| 97 | La Fe | Glioma difuso anaplásico, de fenotipo astrocitario | III | 2 | mut | nd |
| 98 | La Fe | Glioma difuso anaplásico, de fenotipo astrocitario | III | 2 | mut | mut |
| 99 | La Fe | Glioblastoma | IV | 2 | wt | nd |
| 100 | La Fe | Glioma difuso de fenotipo oligodendroglial | II | 1 | mut | wt |
| 101 | La Fe | Oligodendroglioma | II | 1 | mut | wt |
| 102 | La Fe | Glioblastoma | IV | 2 | wt | wt |

### Supplementary Table 2. Paired human samples

| Sample | Hospital | Year of surgery | Progression Free Survival | Diagnosis | Grade | Tau IHC | IDH | ATRX |
|---|---|---|---|---|---|---|---|---|
| 3 | H12O | 2010 | | Astrocytoma | III | 2.6 | mut | mut |
| | H12O | 2011 | 18 months | Astrocytoma | III | 1.6 | mut | mut |
| 5 | H12O | 2009 | | Astrocytoma | II | 2.3 | mut | mut |
| | H12O | 2011 | 22 months | Glioblastoma | IV | 0.7 | mut | mut |
| 6 | H12O | 2008 | | Oligodendroglioma | III | 1.6 | mut | wt |
| | H12O | 2011 | 29 months | Oligodendroglioma | III | 1.5 | mut | wt |
| 7 | H12O | 2014 | | Astrocytoma | II | 1.2 | mut | mut |
| | H12O | 2016 | 19 months | Glioblastoma | IV | 0.2 | mut | mut |
| 8 | H12O | 2013 | | Astrocytoma | II | 1.7 | mut | mut |
| | H12O | 2018 | 50 months | Astrocytoma | III | 0.8 | mut | mut |
| 10 | H12O | 2010 | | Astrocytoma | II | 1.2 | mut | mut |
| | H12O | 2018 | 100 months | Astrocytoma | II | 1.7 | mut | mut |
| 11 | H12O | 2012 | | Astrocytoma | III | 1.2 | mut | mut |
| | H12O | 2015 | 39 months | Glioblastoma | IV | 0.0 | mut | mut |
| 13 | H12O | 2013 | | Astrocytoma | II | 1.5 | mut | mut |
| | H12O | 2016 | 42 months | Astrocytoma | II | 2.6 | mut | mut |

**Supplementary Table 3. GB cell lines.** (nd: not diagnosed. 0: wild type. 1: altered)

| Cell line | Origin | EGFR amp | EGFR mut | PTEN loss | p53 mutation |
|---|---|---|---|---|---|
| **RG1** | Mazzoleni et al. | 1 | 0 | 0 | 1 |
| **12o01** | Hospital 12 de Octubre | 1 | 1 (vIII) | 0 | 1 |
| **12o02** | Hospital 12 de Octubre | 0 | 0 | 1 | 1 |
| **12o12** | Hospital 12 de Octubre | 1 | 1(vIII) | nd | 0 |
| **12o15** | Hospital 12 de Octubre | 0 | 0 | 0 | 1 |
| **12o16** | Hospital 12 de Octubre | 0 | mut | nd | nd |
| **12o22** | Hospital 12 de Octubre | 1 | 1(vIII) | nd | nd |
| **12o29** | Hospital 12 de Octubre | nd | nd | nd | nd |
| **12o33** | Hospital 12 de octubre | 1 | nd | nd | nd |
| **12o40** | Hospital 12 de octubre | nd | nd | nd | nd |
| **12o43** | Hospital 12 de Octubre | nd | nd | nd | nd |
| **12o44** | Hospital 12 de Octubre | nd | nd | nd | nd |
| **12o49** | Hospital 12 de Octubre | 1 | 1 (vII) | nd | nd |
| **12o53** | Hospital 12 de Octubre | nd | nd | nd | nd |
| **12o56** | Hospital 12 de Octubre | nd | nd | nd | nd |
| **12o84** | Hospital 12 de Octubre | 1 | 1<br>C.1118C>A | 0 | 0 |
| **12o89** | Hospital 12 de Octubre | 0 | 1<br>V774M | mut | 1 |
| **12o107** | Hospital 12 de Octubre | 0 | 0 | 0 | 1 |
| **12o108** | Hospital 12 de Octubre | 0 | 0 | 0 | 0 |
| **12o113** | Hospital 12 de Octubre | 1 | 0 | 0 | 0 |
| **12o116** | Hospital 12 de Octubre | nd | nd | nd | Nd |
| **12o121** | Hospital 12 de Octubre | 1 | 0 | 1 | 0 |
| **12o124** | Hospital 12 de Octubre | 1 | nd | nd | nd |
| **12o126** | Hospital 12 de Octubre | 0<br>1 | 1<br>G1134S | 1 | 1 |
| **12o129** | Hospital 12 de Octubre | Nd | nd | nd | nd |
| **12o150** | Hospital 12 de Octubre | 0 | 1<br>L760P/M945I | mut | 1 |
| **GB4** | Hospital Ramón y Cajal | nd | nd | Nd | 1 |
| **GB19** | Hospital Ramón y Cajal | nd | nd | nd | 1 |

### Supplemetary Table 4. Antibodies

| Antibody | Dilution | Source |
|---|---|---|
| **Acetyl-Tubulin** | 1:5000 (WB) | Sigma |
| **AKT** | 1:1000 (WB) | Cell Signaling (4691) |
| **ANGPT2** | 1:100(IF) | Santa Cruz Biotechnology (SC-74403) |
| **α SMA** | 1:500 (WB) 1:100 (IHC) | Santa Cruz Biotechnology (SC-32251) |
| **β-Actin** | 1:1000 (WB) | Sigma |
| **β-catenin** | 1:1000 (WB) | Cell |
| **BrdU** | 1:100 (IHC) | Dako |
| **CD248** | 1:500 (WB) | Santa Cruz Biotechnology (SC-377221) |
| **EGFR** | 1:1000 (WB) | Cell Signaling (2232) |
| **Endomucin** | 1:100 (IF) | Santa Cruz Biotechnology (SC-65495) |
| **GAPDH** | 1:500 (WB) | Santa Cruz Biotechnology (SC-47724) |
| **GFP** | 1:100(IHC) | Santa Cruz Biotechnology |
| **NG2** | | |
| **OLIG2** | 1:100(IHC) | Santa Cruz Biotechnology |
| **phospho-Ser431 AKT** | 1:1000 (WB) | Cell Signaling (4549) |
| **phospho-Tyr751 PDGFRB** | 1:1000 (WB) | Cell Signaling (4549) |
| **phospho-Tyr740 PDGFRB** | 1:100 (IHC) | SIGMA (SAB4504202) |
| **PDGFRB** | 1:1000 (WB) | Cell Signaling (4564) |
| **phospo-Tyr1068 EGFR** | 1:1000 (WB) | Cell Signailing (3777) |
| **phospo-Ser536 NFK β** | 1:1000 (WB) | Cell Signaling (3033) |
| **TAU 7.51** | 1:500 (WB) | |
| **TAU-5** | 1:500 | Calbiochem (577801) |
| **TAU-12** | 1:500 | Millipore (MAB2241) |
| **TAU** | (IHQ) | Dako |
| **YAP-TAZ** | 1:1000 (WB)<br>1:50 (IHC) | Cell Signaling<br>SIGMA |

| | | |
|---|---|---|
| **anti mouse -Dylight 488** | 1:500 (IF) | Jackon Immunoresearch |
| **anti rabbit -Dylight 488** | 1:500 (IF) | Jackon Immunoresearch |
| **Anti mouse-Cy3** | 1:500 (IF) | Jackon Immunoresearch |
| **Anti rabbit-Cy3** | 1:500 (IF) | Jackon Immunoresearch |
| **Anti rat-Cy5** | 1:500 (IF) | Jackon Immunoresearch |

**Supplementary Table 5. qRT-PCR primers**

| Specie | gene | Forward (5'-3´) | Reverse (3'-5´) |
|---|---|---|---|
| mouse | CD31 | TCCAGGTGTGCGAAATGCT | TGGCAGCTGATGCCTATGG |
| mouse | ENG | TGCACTTGGCCTACGACTC | TGGAGGTAAGGGATGGTAGCA |
| mouse | VE-CAD | TTACTCAATCCACATACACATTTTCG | GCATGATGCTGTACTTGGTCATC |
| mouse | CD248 | TTGATGGCACCTGGACAGAGGA | TCCAGGTGCAATCTCTGAGGCT |
| mouse | αSMA | ACCATCGGCAATGAGCGTTTCC | GCTGTTGTAGGTGGTCTCATGG |
| mouse | PDGFRB | CCGGAACAAACACACCTTCT | TATCCATGTAGCCACCGTCA |
| mouse | MMP9 | GCAAGGGGCCGTGTCTGGAGATTC | GCCCACGTCGTCCACCTGGTT |
| mouse | LMNA | TTGCCTCAACTGCAATGACAA | TCTCGATGTCGGTAAAACCCC |
| mouse | KDR | TTTGGCAAATACAACCCTTCAGA | GCAGAAGATACTGTCACCACC |
| mouse | VEGFR2 | CATCACCGAGAACAAGAACAAAACT | GATACCTAGCGCAAAGAGACACATT |
| mouse | TEK | ACGGACCATGAAGATGCGTCAACA | TCACATCTCCGAACAATCAGCCTGG |
| mouse | NRP1 | GCTTGTGCTCTATGCAGATCG | TCGACGAACTCCTGGTGATTTA |
| mouse | EPHA2 | GCACAGGGAAAGGAAGTTGTT | CATGTAGATAGGCATGTCGTCC |
| mouse | AQP1 | AGGCTTCAATTACCCACTGGA | GTGAGCACCGCTGATGTGA |
| mouse | VEGF A | TGCCAAGTGGTCCCAGGCTGC | CCTGCACAGCGCATCAGCGG |
| mouse | PDGF A | GATACCTCGCCCATGTTCTG | CAGGCTGGTGTCCAAAGAAT |
| mouse | PDGF B | GGGCCCGGAGTCGGCATGAA | AGCTCAGCCCCATCTTCATCTTACGG |
| mouse | PGF | GAGGCCAGAAAGTCAGGGGGC | ATGGGCCGACAGTAGCTGCGA |
| mouse | CCL2 | AGGTCCCCTGTCATGCTTCTG | TCTCCAGCCTACTCATTGGG |
| mouse | PI3KCG | GCTCTTCGCCAATCACACAAAC | GGCATTCCTGTCATCAGCATC |
| mouse | NG2 | GACGGCGCACACACTTCTC | TGTTGTGATGGGCTTGTCAT |
| mouse | Ang1 | CATTCTTCGCTGCCATTCTG | GCACATTGCCCATGTTGAATC |
| mouse | Ang2 | TTAGCACAAAGGATTCGGACAAT | TTTTGTGGGTAGTACTGTCCATTCA |
| human | MAPT | GTCGAAGATTGGGTCCCTGG | GACACCACTGGCGACTTGTA |
| human | αSMA | TAGCACCCAGCACCATGAAGATCA | GAAGCATTTGCGGTGGACAATGGA |
| human | NG2 | AGCTCTACTCTGGACGCC | ATCGACTGACAACGTGGC |
| human | CD248 | AGACCACCACTCATTTGCCTGGAA | AGTTGGGATAATGGGAAGCGTGGT |
| human | PDGFRB | ACGGCTCTACATCTTTGTGCCAGA | TCGGCATGGAATGGTGATCTCAGT |
| human | SNAIL 1 | ACCACTATGCCGCGCTCTT | GGTCGTAGGGCTGCTGGAA |
| human | SNAIL 2 | ATCTGCGGCAAGGCGTTTTCCA | GAGCCCTCAGATTTGACCTGTC |
| human | ZEB 1 | GGCATACACCTACTCAATACGG | TGGGCGGTGTAGAATCAGAGTC |
| human | ZEB 2 | AATGCACAGAGTGTGGCAAGGC | CTGCTGATGTGCGAACTGTAGC |
| human | TWIST 1 | CCGGAGACCTAGATGTCATTG | CACGCCCTGTTTCTTTGAAT |
| human | SERPINE | CATAGTGGAAGTGATAGAT | ACTCTGTTAATTCGTCTT |
| human | TAZ | TTTCCTCAATGGAGGGCCA | GGGTGTTTGTCCTGCGTTTT |
| human | CD133 | GCCACCGCTCTAGATACTGC | TGTTGTGATGGGCTTGTCAT |
| human | SOX2 | GCGAACCATCTCTGTGGTC | AATGGAAAGTTGGGATCGAA |
| human | L1 CAM | TCGCCCTATGTCCACTACACCT | ATCCACAGGGTTCTTCTCTGGG |
| human | DLL3 | AAACCTATGGGCTTGAGGAG | CGCTGAGTACAATCAGTGGAA |
| human | REST | CGGCTAACAATACTAATCG | TAGACTCGCTCATTATCC |
| human | LIF | CATGAACCAGATCAGGAG | GCTGTGTAATAGAGAATAAAGAG |
| human | NESTIN | GCGGCTGCGGGCTACTGAAA | CCAGCTGCTGCCGACCTTCC |
| human | OLIG2 | CGGCTTTCCTCTATTTTGGTT | GTTACACGGCAGACGCTACA |

**Supplementary Table 6. Sequencing primers**

| Species | Gene name | Forward (5'-3´) | Reverse (3'-5´) |
|---|---|---|---|
| **human** | **EGFR** | CAGTATTGATCGGGAGAG | CATCTCATAGCTGTCGGC |
| | | CAACATGTCGATGGACTTCCA | TTCGCATGAAGAGGCCGATC |
| | | GCCCCCACTGCGTCAAGACC | AGCTTTGCAGCCCATTTCTA- |
| | | CAGCGCTACCTTGTCATTCA | CTATCCTCCGTGGTCATGCT |
